# Unlocking the Potential of High-Quality Dopamine Transporter Pharmacological Data: Advancing Robust Machine Learning-Based QSAR Modeling

**DOI:** 10.1101/2024.03.06.583803

**Authors:** Kuo Hao Lee, Sung Joon Won, Precious Oyinloye, Lei Shi

## Abstract

The dopamine transporter (DAT) plays a critical role in the central nervous system and has been implicated in numerous psychiatric disorders. The ligand-based approaches are instrumental to decipher the structure-activity relationship (SAR) of DAT ligands, especially the quantitative SAR (QSAR) modeling. By gathering and analyzing data from literature and databases, we systematically assemble a diverse range of ligands binding to DAT, aiming to discern the general features of DAT ligands and uncover the chemical space for potential novel DAT ligand scaffolds. The aggregation of DAT pharmacological activity data, particularly from databases like ChEMBL, provides a foundation for constructing robust QSAR models. The compilation and meticulous filtering of these data, establishing high-quality training datasets with specific divisions of pharmacological assays and data types, along with the application of QSAR modeling, prove to be a promising strategy for navigating the pertinent chemical space. Through a systematic comparison of DAT QSAR models using training datasets from various ChEMBL releases, we underscore the positive impact of enhanced data set quality and increased data set size on the predictive power of DAT QSAR models.

## The dopamine transporter and its inhibitors

Neurotransmitter:sodium symporters (NSSs) transport synaptic neurotransmitters through a Na^+^ and Cl^--^-dependent mechanism back to the presynaptic neuron ^1, 2^. Among NSSs, the monoamine transporters include the dopamine, serotonin, and the norepinephrine transporters (DAT, SERT, and NET, respectively) ^3^. The cognate substrates of these transporters play key roles in the central nervous system (CNS), responsible for mood, emotion, learning, cognition, memory, sleep, and appetite. Inhibition of these transporters leads to reduced clearance of these neurotransmitters, resulting in prolonged synaptic signaling with higher intensity ^3^.

Specifically, dopamine is an essential in the brain’s reward system ^4^. DAT is responsible for regulating the extracellular dopamine levels in the brain, using the energy stored in the Na^+^ and Cl^-^ gradients to symport dopamine back to the neuron ^5, 6^. Disrupting the DAT function can lead to several psychiatric disorders, including attention-deficit hyperactivity disorder (ADHD) ^7, 8^, bipolar disorder ^9, 10^, and depression ^11^. Many psychostimulants, including cocaine and amphetamine, primarily target DAT. Upon entering the human brain, psychostimulants elevate extracellular dopamine levels and can have a highly addictive effect on the individuals consuming the substances, with the possibility of triggering substance use disorders (SUDs).

SUDs encompass complex health conditions that include significant impairments of physiological, mental, and social functions due to consumption of substances at high doses and/or frequencies ^12^. Specifically, cocaine use disorder (CUD) is characterized by the compulsive consumption of cocaine despite its adverse medical, psychological, and behavioral consequences, was found to affect more than 5 million individuals in 2019 ^13^. Notably, cocaine overdose death rates increased nearly 54% from 2019 to 2021 ^14^. While users experience a brief sense of intense euphoria when consuming cocaine before the effect wears off, prolonged usage is linked to the development of mental disorders (e.g., depression) and cognitive impairments ^13^. Although cocaine acts non-selectively on the three monoamine transporters, substantial evidence indicates that DAT plays a key role in the development of CUD ^15, 16^.

Many efforts have been spent in past decades toward developing effective medications for CUD; however, there is no approved FDA drugs for CUD. Early efforts in developing therapeutic agents for CUD have centered around cocaine analogues, which have been optimized to possess both higher affinity and improved selectivity for the DAT but decreased stimulatory effects and less abuse liability ^3, 17, 18^. It was then discovered that the compounds with decreased stimulatory effects stabilized the DAT in a distinct conformation as compared to cocaine ^17, 19^. These compounds were classified as “atypical” DAT inhibitors, distinguishing them from “typical” DAT inhibitors like cocaine ^18–20^. Thus, cocaine stabilizes DAT in an outward-facing conformation, whereas atypical inhibitors, such as modafinil and JHW007, favor inward-facing conformations ^21–23^. Notable examples of atypical DAT inhibitors include benztropine ^17, 24^, rimcazole ^17^, GBR12909 ^25^, and modafinil analogues ^26, 27, 28^.

However, translating these compounds to a human CUD treatment has been challenging ^29^, while the potential for discovering additional candidate atypical DAT inhibitors based on the well-characterized scaffolds is diminishing. In this work, we provide a comprehensive overview from the cheminformatics perspective in identifying novel DAT inhibitor scaffolds.

## Ligand-based versus structure-based drug discovery for the NSSs

Computer-aided drug design (CADD) can be broadly categorized into protein structure-based drug design (SBDD) and ligand-based drug design (LBDD) ^30^. SBDD leverages in-depth knowledge of the three-dimensional (3D) structures of the protein targets, allowing the design of small molecules that can interact optimally with the targets ^31^. When the high-resolution structures of mammalian NSSs were unavailable, SBDD heavily depended on the qualities of homolog-modeling based computational models of the targets, emphasizing the importance of accurate and reliable molecular modeling approaches ^32–36^. LBDD, on the other hand, operates without the 3D structural information of the targets, but relies on the analysis of the known pharmacological information between small molecules (ligands) and their targets. LBDD is particularly useful when the 3D structure of the target is either unknown or challenging to obtain ^37^. Thus, while SBDD provides the target structure-guided precision, LBDD offers ligand-based adaptability, in the quest for new therapeutic agents.

In LBDD, several popular techniques have been employed, with notable examples being pharmacophore modeling and quantitative structure-activity relationship (QSAR) modeling. In pharmacophore modeling, pharmacophore refers to the spatial arrangements of structural and chemical features, usually on a chemical scaffold, that can be extracted from the active compounds of a target. The resulting model can facilitate molecular recognition to effectively identify and characterize new compounds ^38–40^. QSAR modeling was first established by Corwin Hansch for its application in the field of virtual drug screening ^41^. QSAR modeling focuses on establishing a quantitative relationship between the chemical structures of small compounds and their specific pharmacological activities, by building models that correlate the structural features of compounds with their observed bioactivities. The QSAR method operates on the premise that molecules sharing similar structural or physicochemical properties would demonstrate comparable biological activities ^42^. Consequently, the robustness of the QSAR models depends on both the quantity and quality of available pharmacological activity and compound data, much of which have been systematically accumulated and curated in pertinent databases ^43^. These models not only provide valuable insights into the essential molecular features contributing to the desired pharmacological effects, but also aid in the design and optimization of novel drug candidates.

The molecular descriptors that QSAR utilizes come in many different forms, including quantitative (molecular shape) and qualitative descriptors (fingerprints) ^44^, and can range from 2D to 6D ^45^. 2D molecular descriptors, which are the most popular, provide information on the connectivity of atoms, properties of chemical bonds, and chemical fingerprints ^45^. Many commercial software, such as Mold^2^ and DRAGON system, can be used to generate 2D molecular descriptors ^46, 47^. 3D descriptors provide physical information, such as surface properties, molecular volume, and molecular interaction fields ^48–50^. A caveat to 3D descriptors is that these additional complexities may not be necessary to improve the predictive power of QSAR models at the expense of the significantly more computational cost, as its performance is sensitive to the accuracy of predicting ligand conformation.

The early pioneering work in QSAR modeling of DAT ligands includes those of the Newman group, in constructing both 2D and 3D QSAR models, which used 2D and 3D descriptors, respectively. Specifically, the 2D QSAR study was carried out with 70 diverse inhibitors of the DAT and produced robust QSAR models, resulting in a q^2^ = 0.85. These DAT models were then used to search through the National Cancer Institute database, yielding five candidate compounds suitable for testing new DAT scaffolds ^51^. A follow-up study was conducted using 3D QSAR methods for the design of new mazindol analogues and to identify molecular interactions for optimal DAT binding. Using comparative molecular field analysis (CoMFA), each compound’s steric and electrostatic potential fields were calculated and used as features to generate robust 3D QSAR models ^52^.

In recent years, there has been extensive applications of machine learning (ML) and deep learning (DL) techniques in the construction of QSAR models ^53–55^. These advanced computational techniques offer powerful tools for extracting intricate patterns and relationships from complex data sets, beyond the traditional linear regression based QSAR modeling approaches (see below). By leveraging ML and DL algorithms, significantly enhanced accuracy and predictive capabilities of QSAR models have been achieved ^56–60^.

## Databases containing pharmacological activity information of the DAT

Constructing QSAR models involves consideration of many factors, and one of the initial and pivotal aspects is the collection of available and reliable data from relevant databases. We found that the publicly accessible databases containing pharmacological activity information of the DAT include ChEMBL ^61^, DrugBank ^62^, binding database (BindingDB) ^63, 64^, therapeutic target database (TTD) ^65^, psychoactive drug screening program (PDSP) ^66^, and PubChem ^67^. These databases compile data on biomolecules, protein targets, compound characteristics, ADMET properties, binding interactions, functional assays, and more. Typically, these databases are maintained by non-profit organizations and undergo regular updates. While most databases primarily focus on drug-related information and associated targets, BindingDB also offers 3D insights into protein targets and ligands.

To evaluate the available pharmacological data that can be used to construct QSAR models for DAT, we compared pharmacological data of DAT from these databases (data retrieved in December, 2023). Note that in this process, it is essential to consider the number of unique compounds when comparing across databases by removing redundant information. If such information was not found in the database, we employed Morgan fingerprints and Tanimoto similarity measurements to calculate the pairwise similarities to identify the distinct compounds within each database. In addition to human DAT (UniProt ID Q01959), the rattus norvegicus (UniProt ID P23977), bos taurus (UniProt ID P23977), and mus musculus DAT (UniProt ID Q61327) are also included.

- ChEMBL release 33 (May 2023) provides 2,399,743 compounds, 20,334,684 activities, 1,610,596 assays, 15,398 targets, and 88,630 publications ^61^. We found 14,102 pharmacological activity data related to DAT, and among them, we identified 7,496 unique compounds, with clear indications of being curated by experts.
- DrugBank Online (version 5.1.10, released 2023-01-04) contains 15,325 drug entries of FDA-approved drugs or experimental drugs going through the FDA approval process ^62^. Upon querying for the DAT information in DrugBank, our search revealed 51 drugs specifically targeting DAT. Nevertheless, our comparison of data from other databases and literature suggests that additional drugs listed by DrugBank may also bind to DAT (see **Table S1**).
- BindingDB collects experimental data of protein-small molecule interaction ^63, 64^. It provides both 2D- and 3D-information of proteins and ligands. Since its initial launch in the year 2000, BindingDB has accumulated 2.8 million binding data for more than nine thousand targets and over 1.2 million compounds (released 2023-11-30). Regarding DAT, we found 10,650 pharmacological data points, encompassing 5,724 unique compounds.
- TTD provides information of known therapeutic protein targets and the corresponding drugs ^65^. The latest update (2024) includes 3,730 targets and 39,862 drugs. We found 3,578 pharmacological data points directed at DAT. Within that data set, some compound structures are unavailable, making it a challenge to identify unique compounds directly.
- PDSP provides data assessing pharmacological and functional effects at CNS receptors, channels, and transporters ^66^. We found 1,234 instances of pharmacological data associated with DAT, which are related to 327 unique compounds.
- PubChem is one of the largest chemical databases of publicly accessible chemicals ^67^. There are hundreds of data sources connected to PubChem, containing over 116M compounds, 309M substances, 229M bioactivities, 36M publications and 38M patterns. When focusing on DAT specifically, we found a total of 9,108 pharmacological activities, related to 6,021 distinct compounds. However, the level of expert curation on these DAT activities are not clear.

Among these six databases, ChEMBL is clearly outstanding, as it amasses the largest number of unique compounds and offers an extensive repository of expert-curated bioactivity information. Consequently, it was our primary choice for sieving training data for the development of a robust DAT QSAR model ^68^. However, we note that BindingDB is another important resource for DAT pharmacological activity data.

To illustrate the trend of the evolution and the accumulation of DAT pharmacological data over the past four decades, we queried ChEMBL 33 and compiled the retrieved DAT pharmacological activity data along the years (**Fig. 1**, see its legend for query criteria). The first publication curated by ChEMBL on DAT studied amphetamine analogs and tested their inhibition of dopamine uptake in rats ^69^. Along the years, the number of relevant publications as well as the pharmacological activity data have been accumulating substantially (**Fig. 1A,B**). In particular, the number of publications had a large spike in 2008 (55 publications), possibly attributed to the availability of the crystal structure of a bacterial NSS homolog, the leucine transporter (LeuT) in 2005 ^70^), which allowed homolog modeling of the DAT and other NSSs with a high-resolution experimentally determined structure as the template. Integrated with molecular dynamics simulations and uptake experiments, the LeuT-based homology DAT models have been used to understand the substrate binding mode and the molecular mechanism of inhibitor interactions with DAT ^71–74^. Importantly, the LeuT-based homology DAT models have provided valuable guidance to facilitate the drug discovery ^36, 75^. However, we did not detect that the publication of the drosophila DAT structure in 2013 ^76^ resulted in another obvious large accumulation in the relevant publications, although the pharmacological activity data and the number of new unique compounds somewhat increased in 2014 as compared to those in 2013 (**Fig. 1B,C**).

**Figure 1.**
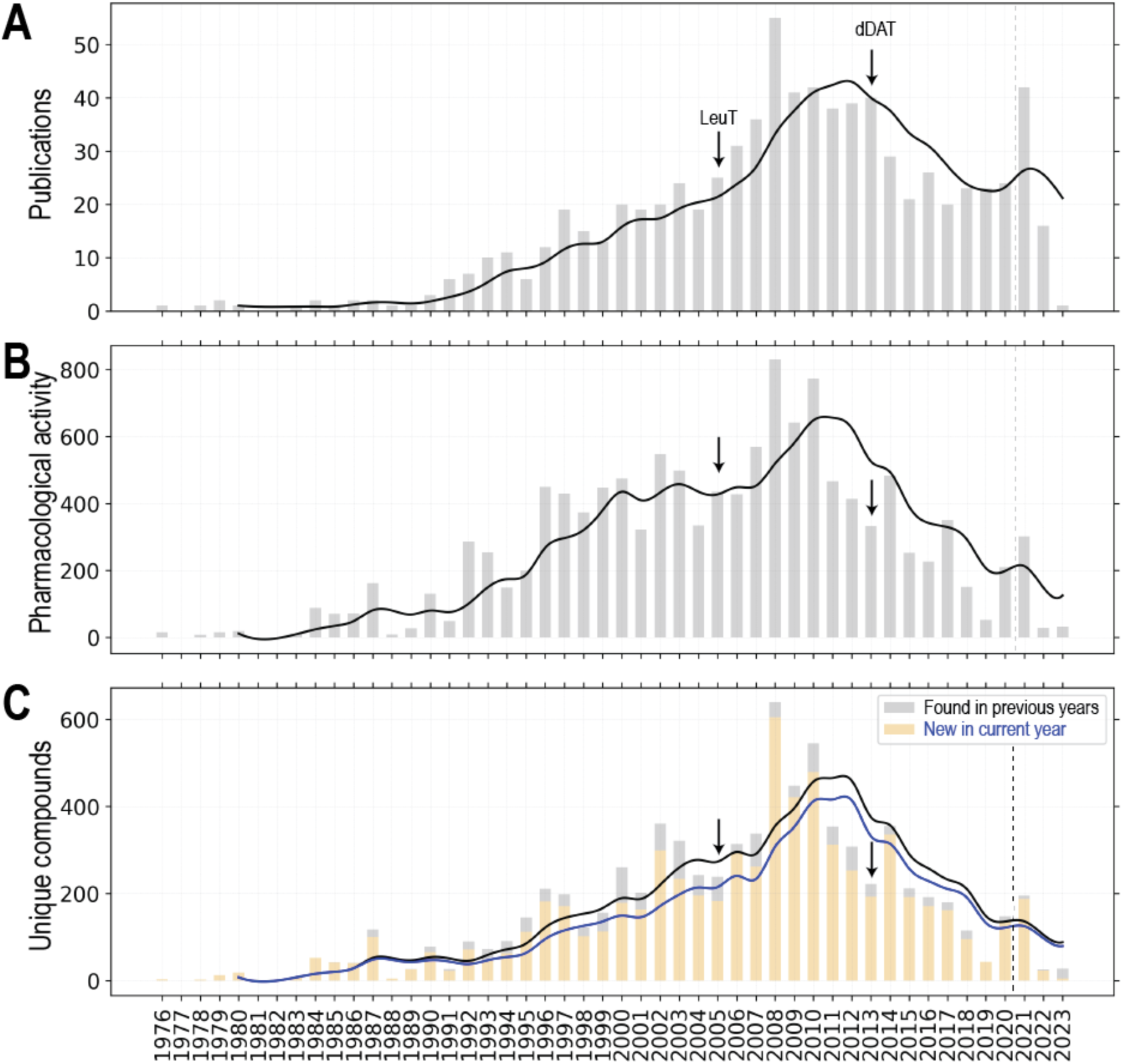
Statistics of DAT activity data from ChEMBL33. The DAT data set was queried and retrieved from entries in a locally installed instance of ChEMBL 33 (May 2023 release). We employed the same query criteria as in our previous study to filter the DAT pharmacological data set ^68^. Number of publications on DAT ligands (A), number of related pharmacological activity data (B) and number of unique DAT ligands (C) were plotted along the years. If a compound has <0.999 pairwise similarity to all the other compound in the data set, we consider it as a unique compound. For panel C, the new unique compound in the current year is colored wheat, and the unique compounds found in previous years is in light grey. The total number of publications, pharmacological activity data, and unique compounds are 773, 7815, and 6689 respectively. The data sets in each panel are separated by dash lines between 2020 and 2021, to indicated what we observed that the curation in ChEMBL can be delayed for more than 2 years.

Notably, whereas the number of publications appears to be plateaued in recent years (**Fig. 1A**), between 2008-2010 and 2019-2020, there has been an obvious decline of novel DAT inhibitors being discovered and the associated pharmacological characterizations (**Fig. 1B,C**). Even though there is a noticeable increase of related publications in 2021, the increase of novel DAT ligands was minimal (note that to indicate the delay of ChEMBL data curation, which we found could be more than two years, a dashed line was added to separate the data of year 2021-2023 from early years in **Fig. 1**).

## Some of representative DAT ligands specifically and not specifically developed for DAT

When targeting a specific protein, such as DAT, for therapeutic purpose, it is critical the understand the pharmacology beyond the target. Conversely, some high-affinity DAT ligands were not necessarily specifically developed for DAT. To evaluate whether the SAR information accumulated by the field focusing on DAT ligand development can be supplemented by other efforts, we gathered and analyzed a set of representative DAT ligands from both the well-respected review articles on the DAT inhibitors ^21, 22, 77^ and DrugBank ^62^, which collects and curates the compounds that have progressed significantly along the drug development processes. We identified those with relatively high affinities using dopamine as a reference (with either pKi, pIC50, pEC50, or pKd > 5.06) to narrow down to a total of 61 ligands (**Table S1**).

Among the 61 ligands, 48 ligands can be found in ChEMBL, but only 46 have recorded DAT pharmacological activity therein. In addition to ChEMBL, 52 ligands can be found in BindingDB, 38 ligands can be found in PDSP, emphasizing the necessity to source the data beyond an individual database. To collect adequate data for **Table S1**, we also had to search the literature extensively.

To characterize the chemical features of these 61 representative DAT ligands, we clustered them based on their chemical structures (see **Fig. 2** legend for the clustering algorithm we employed). Cocaine and eight other DAT ligands were clustered together, and share both a tropane and a phenyl ring. However, a noticeable difference between cocaine and other compounds is the linkage distance between the tropane and phenyl ring. For cocaine, they are separated by a carboxyl group, but for others, they are directly connected. Interestingly, cocaine has the least binding affinity compared to the other compounds in this cluster. Indeed, compared to the affinities at SERT and NET, cocaine is not a DAT selective ligand, while RTI-55, RTI-82, MFZ 2-24, ioflupane I-123 and WIN 35,428 have been used as radiolabeled probes in the binding assays for DAT ^78–82^. Among them, we could not found activity data at SERT or NET for RTI-82, MFZ 2-24 and ioflupane I-123, suggesting that these cocaine analogues may have been primary designed for DAT. Altropane is the only DAT selective ligand in this cluster, and has been used as an imaging probe to monitor DAT ^83^. Troparil (WIN 35,065-2) is a common inhibitor for the three monoamine transporters, and has exhibited a stronger affinity at DAT compared to that of cocaine ^19, 84^. Tesofensine, another common inhibitor for the three transporters, was investigated for its potential applications in obesity-related conditions, Parkinson’s disease, and Alzheimer’s disease ^85^.

**Figure 2.**
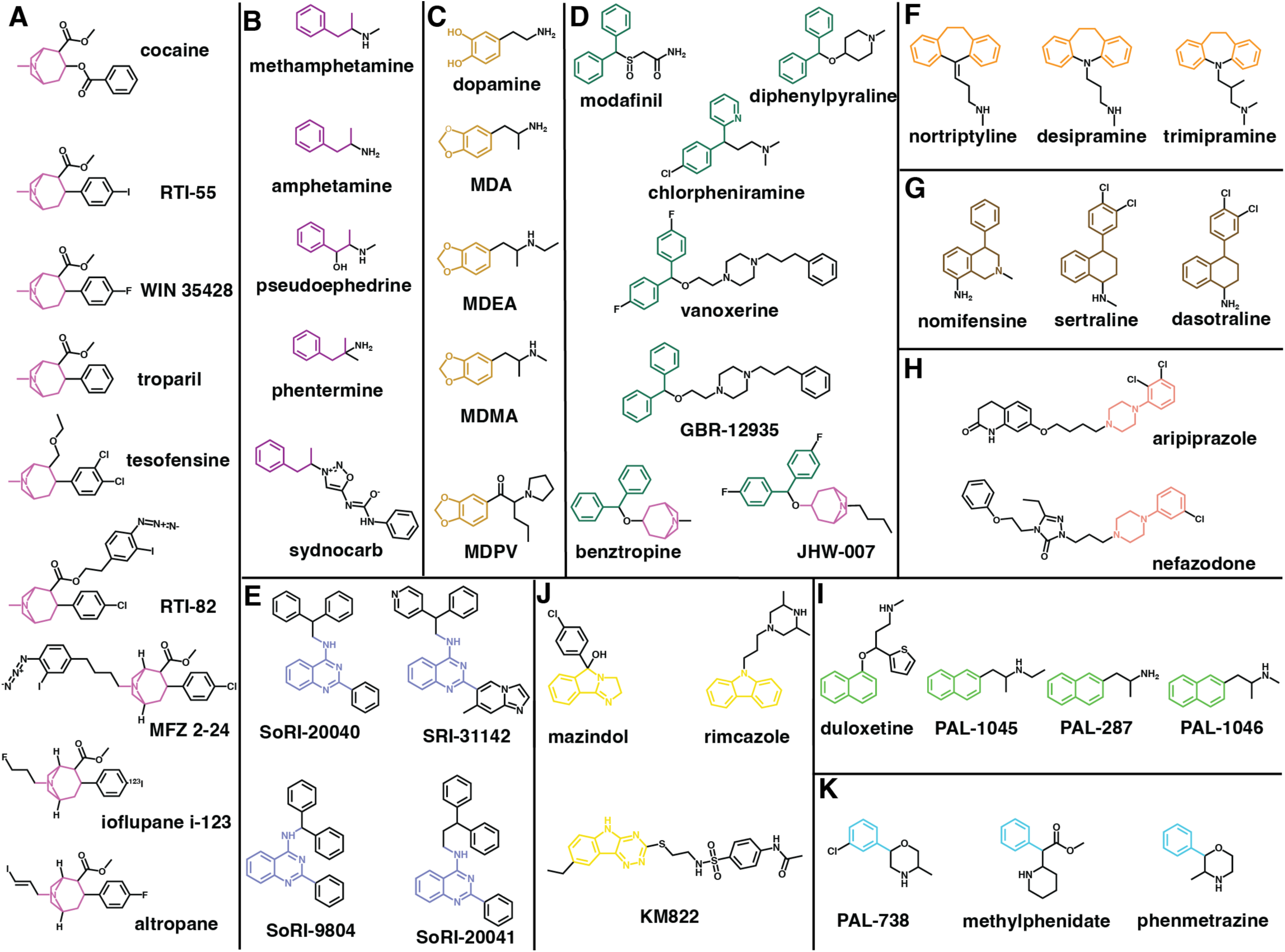
Clustering of representative DAT ligands. For a total of 61 well-known DAT inhibitors collected from literature ^21, 22, 77^ and DrugBank ^62^. To characterize these DAT ligands, we employed the hierarchical clustering approach implemented in the Schrodinger Maestro suite (version 2023-3). We used linear fingerprints, Tanimoto similarity, and average linkage method. Note that three pairs of compounds (armodafinil and modafinil, methylphenidate and dexmethylphenidate, and amphetamine and dextroamphetamine) are enantiomers and we only use modafinil, methylphenidate, and amphetamine when carrying out the clustering. Some known scaffolds were also used to adjust the final representative clusters. The single-member clusters are shown in **Fig. S1**.

In the cluster containing methamphetamine, amphetamine, and three other ligands, the common feature among them is a propylbenzene scaffold linked to a protonated amine (**Fig. 2B**). Note that most ligands in this cluster are DAT substrates except for sydnocarb.

Amphetamine has been used medically to aid in treating ADHD ^86, 87^ and in enhancing cognitive functions ^88^. Due to the size and structure of amphetamine analogs, they can cross through the blood brain barrier (BBB) ^89^. However, it has been found that methamphetamine disrupts the BBB and affects its structural integrity and permeability ^90^. Methamphetamine is a recreational psychostimulant drug known for its strong neurotoxic and addictive effects ^91^. Methamphetamine use disorder (MUD) exhibits a wide spectrum of adverse outcomes from hallucinatory ideation to self-injurious behaviors ^92^. Notably, MUD has been linked to severe cardiac complications ^92^. Both methamphetamine and amphetamine function as potent releasers that increase the extracellular concentrations of key neurotransmitters such as dopamine and serotonin in the brain ^92, 93^. Sydnocarb functions as a noncompetitive inhibitor of DAT and is used for the treatment of schizophrenia and depression but has been linked to having psychostimulant effects ^94, 95^.

Dopamine is an endogenous substrate of DAT. It can be clustered together with several benzodioxole compounds, including 3,4-methylenedioxymethamphetamine (MDMA), 3,4-methylenedioxypyrovalerone (MDPV), 3,4-methylenedioxy-N-ethylamphetamine (MDEA), and 3,4-methylenedioxyamphetamine (MDA) (**Fig. 2C**). MDMA, commonly known as “ecstasy”, acts on both DAT and SERT, and can have stimulatory effects and alter perception and induces hallucinatory experiences ^96–98^. MDEA, functions as a partial releaser of DAT ^99^, while MDA has been shown to act as a substrate for all three monoamine NSSs ^100^. MDPV, a synthetic cathinone typically seen in “bath salts”, is a psychostimulant drug characterized by its heightened selectivity for DAT and increased potency when compared to that of cocaine ^101^. Studies have shown that MDPV is a DAT inhibitor but not a substrate ^102^, whereas all the other members of this cluster showed at least some substrate properties for DAT ^100, 103^.

A few well-recognized atypical inhibitors were clustered together including benztropine, JHW-007, vanoxerine, modafinil, GBR-12935, diphenylpyraline, and chlorpheniramine (**Fig. 2D**). In this cluster, diphenylmethane is the common moiety. Among them, modafinil is a therapeutic drug involved in treating excessive sleepiness in narcolepsy patients by increasing extracellular levels of dopamine ^27, 104^. Modafinil has two enantiomers (S- and R-modafinil), and it has been shown that the R-enantiomer (also known as armodafinil) has ∼3-fold higher affinity than S-enantiomer at DAT ^27^, while the R-enantiomer is a DAT selective ligand, when comparing the affinities at DAT, SERT and NET ^105^. Vanoxerine (GBR12909), a piperazine derivative and another atypical DAT inhibitor, was initially developed as a therapeutic treatment in combatting cocaine addiction, but it was later discontinued due to its heart-related side effects ^25, 106^. Diphenylpyraline functions as an antihistamine but has also been shown to inhibit DAT. Previous research suggested a potential for it to reduce the effects of cocaine ^107^. Chlorpheniramine is an antihistamine as well and displayed a high affinity for SERT and a relatively low affinity to DAT ^108^. In addition to the diphenylmethane moiety, JHW-007 and benztropine also possess a tropane ring. Benztropine, formally used as a drug for Parkinson’s disease, share noticeable structural similarities with cocaine. It has been shown to function as an atypical DAT inhibitor and utilized as a pharmacological agent for CUD research ^17, 25, 109^. JHW-007 is also an atypical DAT inhibitor and has been demonstrated to possess reduced psychostimulant effects compared to cocaine ^110^. Interestingly, except for diphenylpyraline and chlorpheniramine, the other five ligands in cluster D are selective for DAT over SERT and DAT based on the data that we could collect (**Table S1**).

Some known allosteric inhibitors of DAT were clustered in one group (**Fig. 2E**). Specifically, SRI-31142, is a 4-quinazolinamine derivative that could inhibit the actions of cocaine; however, further research is needed to fully explore its therapeutic potential ^111^. Three other 4-quinazolinamine analogs, SoRI-9804, SoRI-20040, SoRI-20041, are clustered with SRI-31142, and have been found to be partial inhibitors and allosteric modulators for DAT ^112, 113^.

While clusters A to E mentioned above have garnered more attentions for their interactions with DAT, the remaining clusters of ligands were mainly developed for use as antidepressants, appetite suppressants, and antipsychotics, which, however, have the off-target effects at DAT. In particular, two clusters of DAT inhibitors were obviously developed to treat depression. They have either tricyclics or bicyclic ring moiety. In the cluster F (**Fig. 2F**), desipramine, nortriptyline, and trimipramine are tricyclic antidepressants (TCAs). Both desipramine and nortriptyline are FDA-approved drugs to treat depression, due to their actions at NET and/or SERT ^114^, and have been shown to have an inhibiting effect on DAT as well ^108^. Trimipramine, has been shown to be a DAT inhibitor but is more potent at SERT ^108^. Nomifensine, sertraline, and dasotraline share a bicyclic ring moiety attached to a toluene group (**Fig. 2G**). Sertraline, a selective serotonin reuptake inhibitor (SSRI) that also inhibits DAT, serves as an antidepressant as well ^115^. Dasotraline is a dual DAT and NET inhibitor, which is being studied for its therapeutic potential in treating ADHD ^116^.

Two other clusters also include antidepressants. One cluster shares phenylpiperazine as the common feature (**Fig. 2H**). In this cluster, aripiprazole is an atypical antipsychotic drug that primary target dopamine D2 receptor with a high affinity ^117^, but can also bind to both DAT and SERT, with only slightly higher affinity at SERT ^118, 119^. Nefazodone is an atypical antidepressant and has similar affinities in the three monoamine NSSs ^120^. Duloxetine, PAL-287, PAL-1045, and PAL-1046 share a naphthalene moiety and are clustered together (**Fig. 2I**). Duloxetine, a dual NET and SERT inhibitor, has exhibited the potential to serve as an antidepressant based on previous research findings ^121^, but demonstrated a weaker affinity to DAT ^122^. In addition to duloxetine, the other three compounds in cluster I are all substrates. PAL-287 functions as a releaser for DAT, SERT, and NET, and has been shown to suppress the effects of cocaine ^123, Rothman, 2012 #86^ (**Table S1**). PAL-1045 tends to act as a partial releaser for both DAT and SERT ^99, 124^, whereas PAL-1046 functions as a full substrate for DAT ^99^.

Two clusters include appetite suppressants. Phenmetrazine is an anorectic drug used in mid-1900s to promote the release of dopamine in the brain ^125^ and was clustered together with methylphenidate and PAL-738. All three ligands in this cluster share a toluene moiety either linked to a 3-methylmorpholine moiety or carboxyl group (**Fig. 2K**). However, the phenylacetic acid moiety of methylphenidate, which acts as a typical DAT inhibitor, is common to cocaine, but not to the other two members of this cluster. Although methylphenidate serves as a therapeutic drug for ADHD, it also exhibits psychostimulant effects ^126^. PAL-738, on the other hand, serves as a partial releaser for DAT ^99^. The next cluster includes mazindol, which is another appetite suppressant, and a triple reuptake inhibitor of DAT, NET, and SERT ^127, 128^. It can be clustered together with rimcazole and KM822, and they share a similar tricyclic ring moiety (**Fig. 2J**). Rimcazole has been observed to inhibit the cocaine binding at DAT ^129^. KM822 is a noncompetitive inhibitor of DAT and has been shown to reduce the affinity of cocaine to DAT ^35^.

Ten compounds have distinct scaffolds and are the sole representatives in their own respective clusters in this set of 61 ligands (**Fig. S1**). In particular, bupropion functions as a dual DAT and NET inhibitor and is used for the treatment of ADHD, depression, and smoking cessation ^130^. Tamoxifen functions as both a DAT inhibitor with some atypical property without psychostimulant effect and a selective estrogen receptor modulator, which is typically prescribed for the treatment of estrogen related breast cancer ^131^. While having a high affinity at SERT, ibogaine operates as a competitive inhibitor of DAT as well, and is a psychostimulant under investigation for its potential to treat SUDs ^132^.

In summary, while the potential for leveraging the “traditional” DAT inhibitor scaffolds is diminishing, our analysis indicated that ample chemical space can be explored for developing the DAT inhibitors with desired pharmacological profiles. However, the accumulation and availability of large quantity of DAT pharmacological data present both challenges and opportunities.

## Curation and filtering the DAT pharmacological data sets

In order to assemble high-quality training data sets for QSAR modeling, adequate understanding and careful curation of the raw data retrieved from the databases are essential. One important issue is to differentiate the half maximal inhibitory concentration (IC50) and inhibition constant (Ki) in measuring the binding potencies of the ligand at a protein target.

Whereas Ki can be derived from IC50 based on the Cheng-Prusoff equation, Ki is a more accurate representation of binding affinity than IC50, because IC50 can be influenced by the methods used for measurement ^133^. Based on the Cheng-Prusoff equation, Ki values are always expected to be smaller than the IC50 values. However, if the same experimental approaches are employed, the trends of Ki and IC50 should be similar. Additionally, for transporter proteins, we can separate the data according to the assay type, either the radiolabeled inhibitor binding assay (referred to as “binding” below) or uptake inhibition (referred to as “uptake” below), as they represent inhibitory potencies of related but different biological processes ^68^.

Thus, the DAT pharmacological data retrieved from ChEMBL can be divided into four distinct sets: uptake IC50, uptake Ki, binding IC50, and binding Ki data sets (**Table 1**). Notably, among the four distinct data sets, there are some compounds appeared in more than one set (termed as “overlapping” compounds). By analyzing the trends among these compounds, we validate the points described above. First, in both the uptake pKi versus uptake IC50 and the binding pKi versus binding IC50 plots, we can observe that pKi values have a trend of being larger than IC50s, confirming the deduction from the Cheng-Prusoff equation described above (**Fig. 3C,D**). As expected, there is a strong correlation between uptake and binding pKi values, as well as between uptake and binding IC50 values (**Fig. 3A,B**). However, due to limited available data, it is not conclusive whether uptake pKi or pIC50 values tend to be larger than those of binding (**Fig. 3A,B**).

**Figure 3.**
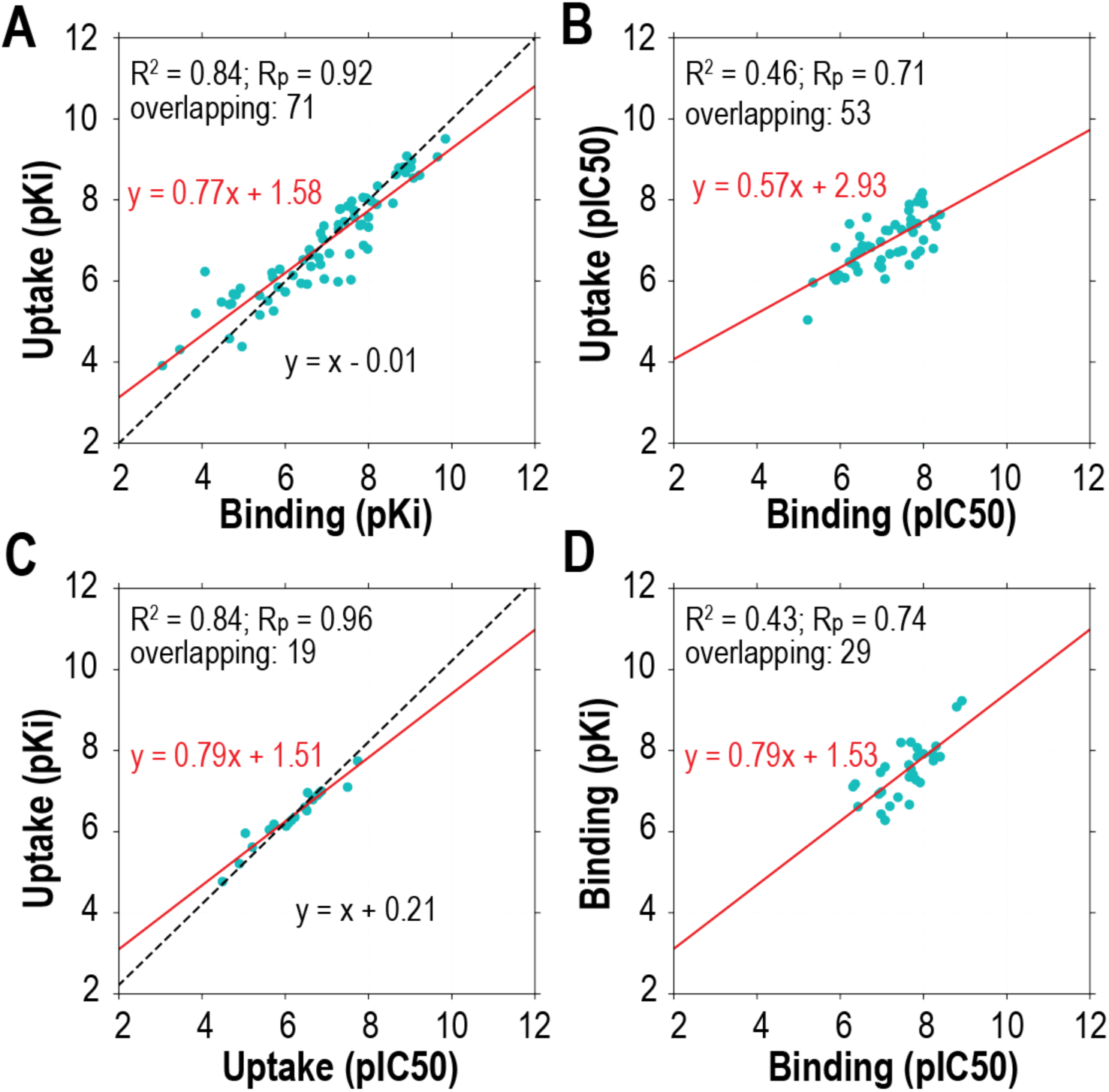
Comparative analysis of DAT pharmacological data sets with correlation metrics and linear regressions. The DAT pharmacological data can be divided into four sets, uptake pKi, uptake pIC50, binding pKi, and binding pIC50. The overlapping compounds between different data sets were extracted for comparisons. The correlation of determination (R^2^) and the Pearson coefficient correlation (Rp), as well as the number of the overlapping compounds (referred as “overlapping”), are indicated at the top left corner of each panel for the indicated comparisons. The red lines are the linear regressions of the indicated data sets; the black dotted lines are the linear regressions with the slope restrained to 1. Note that slope-restrained regression results were not shown for panels B and D, due to poor goodness-of-fit.

**Table 1.** Queried pharmacological activity data and training data sets of the DAT ligands from different ChEMBL releases.

| version | Release date | queried data | uptake |  | binding |  |
| --- | --- | --- | --- | --- | --- | --- |
|  |  |  | Ki | IC50 | Ki | IC50 |
| ChEMBL33 | May 2023 | 14102 (7496) | 366 (366) | 646 (646) | 1166 (1166) | 482 (482) |
| ChEMBL31 | Aug 2022 | 14099 (7470) | 366 (366) | 646 (646) | 1166 (1166) | 482 (482) |
| ChEMBL29 | Jul 2021 | 13915 (7326) | 366 (366) | 646 (646) | 1131 (1131) | 482 (482) |
| ChEMBL27 | May 2020 | 13456 (7049) | 366 (366) | 606 (606) | 1113 (1113) | 482 (482) |
| ChEMBL25 | Mar 2019 | 13273 (6935) | 350 (350) | 593 (593) | 1189 (1189) | 556 (556) |

The numbers of the unique compounds for each data set are shown in the brackets. Note that although some ChEMBL releases have the same numbers for the final training data sets, the value of pharmacological activity data may be different due to the measurement of the same compounds from new studies collected in new ChEMBL releases.

Hence, it is not advisable to combine either Ki and IC50, or uptake and binding data sets in building the QSAR models of DAT and likely other transporter proteins. As there are more Ki data than IC50 data in the binding data sets, we mainly focus on using the binding Ki data for DAT QSAR modeling.

## ML-based QSAR models using DAT binding data set

Machine learning (ML) and deep learning (DL) have been extensively applied in biological and pharmaceutical research, including protein structure prediction ^134, 135^, absorption, distribution, metabolism, excretion, and toxicity (ADMET) profiling ^136, 137^, QSAR modeling ^58, 68^, de novo drug design ^138–140^, and the blood–brain barrier (BBB) permeability prediction ^141^. With the accumulation of large amount of pharmacological data in the databases such as ChEMBL and the drastic improvement of computation hardware in the recent decade, especially the applications of graphics processing units (GPUs) in scientific computing, ML and DL have risen to become the primary approaches in QSAR modeling, by handling the big data sets as well as the high volume of chemical features that can be generated for each compound.

Among many ML algorithms, XGBoost and RF have gained increasing popularity for their strong prediction performances, relative ease in usage, robustness with adjustable hyperparameters, and interpretability ^59, 60, 142^. A comparison of ML- and DL-based QSAR models in the human *ether-à-go-go*-related gene (hERG) indicates that XGBoost provides the best prediction results ^143^. XGBoost was found to be an excellent choice for both large and small data sets for QSAR modeling ^144^. When utilizing the DAT binding data set, XGBoost-trained models exhibited superior performance ^68^.

To evaluate impact of the additional data on the quality of the QSAR models, we performed a benchmark analysis of the DAT binding Ki data sets retrieved from different ChEMBL releases. Across the ChEMBL releases from the past five years, we observed a noticeable enhancement in the coefficient of determination (R^2^) value from 0.72 to 0.77 (**Table 2**). Similarly, mean square error (MSE), Pearson correlation coefficient, and Spearman’s rank correlation coefficient followed this positive trend, with the latest ChEMBL release yielding the most favorable benchmarks.

**Table 2.**
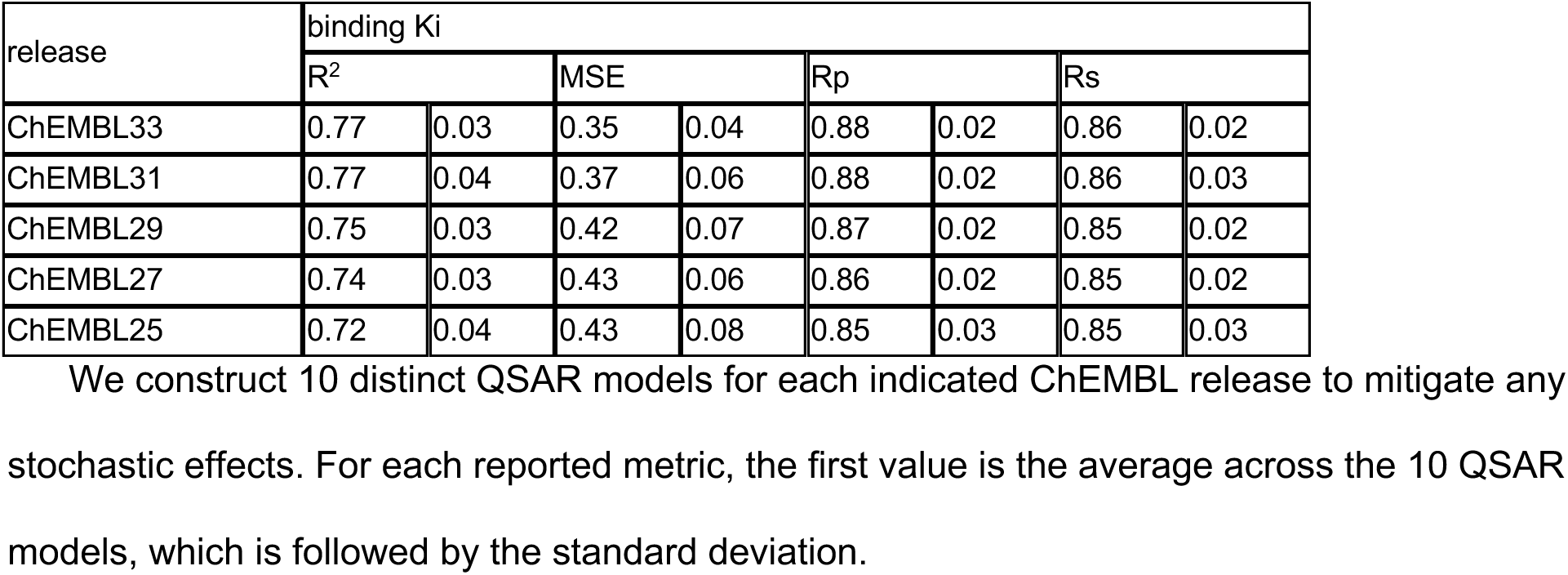
Benchmarks of the DAT QSAR models trained with the XGBoost algorithm.

Note that while overall volume of the queried data continued to grow from ChEMBL 25 to ChEMBL 33, there was no significant increase in the number of unique compounds within the final training data sets (**Table 1**), which may partially be due to the downward trend of medicinal chemistry efforts on DAT (**Fig. 1**), and potentially on NSSs in general. Notably, there were changes of curation criteria between ChEMBL 25 and ChEMBL 27, resulting in a reduction in the number of inhibitor’s Ki and IC50 data sets. These changes in ChEMBL led to an enhancement in the data set’s quality, resulting in a 0.02 increase in the R^2^ value (**Table 2**). Even though there was only a modest increase of 53 data points in the binding Ki training data set from ChEMBL 27 to ChEMBL 33, this change contributed to an overall benchmark improvement of 0.03 in R^2^. Thus, even relatively moderate addition, update, and refinement of data may play a role in enhancing the quality of QSAR models.

While extensive ML/DL efforts have been made on various aspects of the NSSs ^43, 145, 146^, to our knowledge, no DL-based DAT QSAR research has been published. Because proper applications of DL may require larger data sets, we speculate that there are still not yet enough data points available in the public database to build robust DL-based DAT QSAR models. Indeed, ML-based models can use a small training data to still produce robust predictive QSAR models ^147^ and have been seen to outperform DL-based models in various cases ^148^.

## Potential applications and challenge

Virtual screening provides a rapid and low-cost compounds screening in the early stage of drug discovery, by searching potential hits from databases ^149, 150^. Such techniques have been applied in identifying new compounds in SERT ^151–153^. The application of the DAT QSAR model in virtual screening may similarly explore a large compound library and to identify hit compounds and novel scaffolds. In addition, in a medicinal chemistry synthesis campaign, the predicted Ki of candidate compounds by robust DAT QSAR models can help to prioritize the candidate compounds to be synthesized. The iterative prediction, synthesis, and pharmacological measurements are expected to both improve the quality of the QSAR models and improve the efficiency of synthesis campaign.

The screening process can also involve conducting counter- or synergistic-screening with other targets. In particular, we have established the protocol to screen the ligands for the DAT but against the human ether-a-go-go-related gene (hERG) ^68^. The increasing concerns regarding the risks associated with hERG binding have led to a notable surge in the availability of hERG binding affinity data in both public databases and scientific literature. The prediction by the QSAR models can be instrumental in enabling the identification of potential medication candidates characterized by specific attributes, e.g., to aid in pinpointing high-affinity DAT inhibitors with minimal hERG affinity.

At physiological pH of 7.4, dopamine exists predominantly in its cationic form ^154^. pH-induced changes between dopamine and DAT could alter their interactions and conformations. Specifically, the protonation of the amine group not only influences the conformation of dopamine but also introduces a positive charge, enhancing the interaction between the bound dopamine and negatively charged residues in DAT, such as Asp79 in the central binding pocket ^155^. In comparison, modafinil is not in a charged form when it is bound in DAT ^27^. Therefore, considering the proper protonation state of the ligands accordingly should improve the quality of the training data set, by providing a critical chemical context for the models to consider when making bioactivity predictions. However, while different algorithms and software packages are available to predict pKa, such as Epik ^156^ and MolGpka ^157^, which may generate decent pKa predictions in water, it is generally a challenging problem to predict the pKa of the ligands in the binding pocket of the target protein.

## Conclusion

DAT is a key therapeutic target of SUDs. By collecting and analyzing the representative DAT ligands, we reveal that DAT binds to a variety of compounds, while there is still a large chemical space to be explored for novel DAT ligand scaffolds. The compilation of extensive DAT pharmacological activity data sets, coupled with the power of QSAR modeling, presents a promising avenue for exploring such a space.

## Acknowledgements

Support for this research was provided by the National Institute on Drug Abuse–Intramural Research Program, Z1A DA000606 (L.S.).

## Supplemental Materials

**Table S1.**
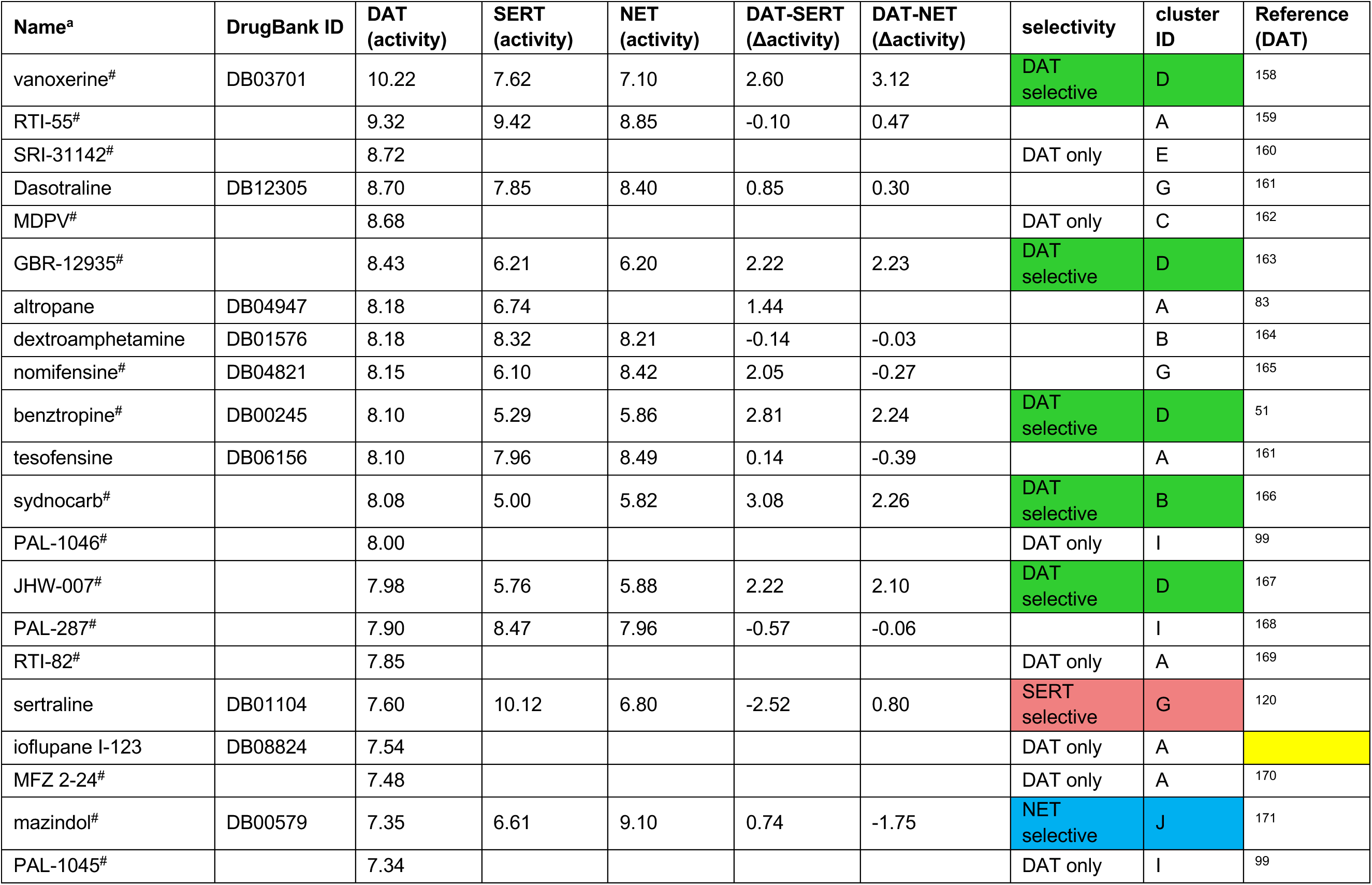
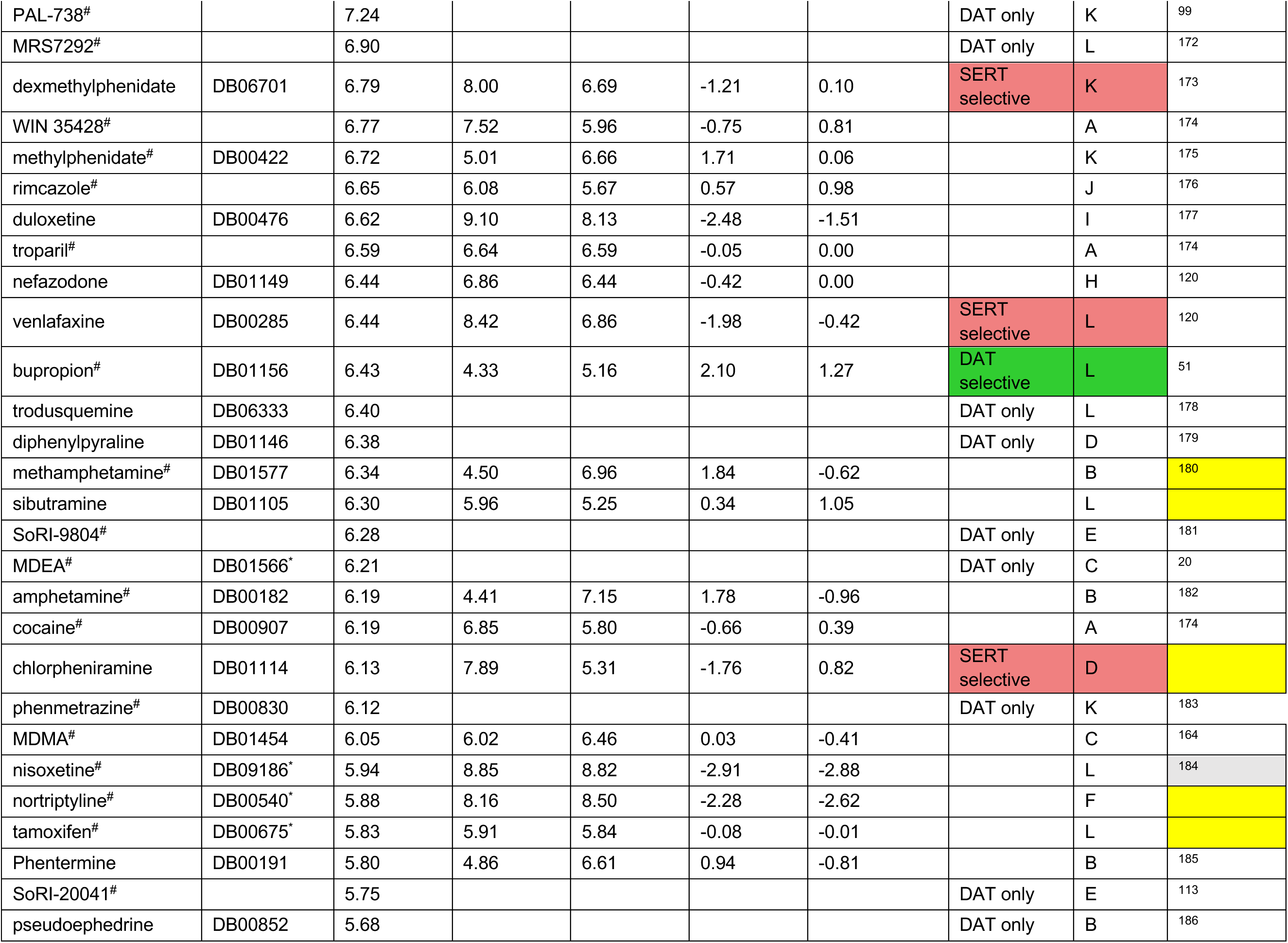
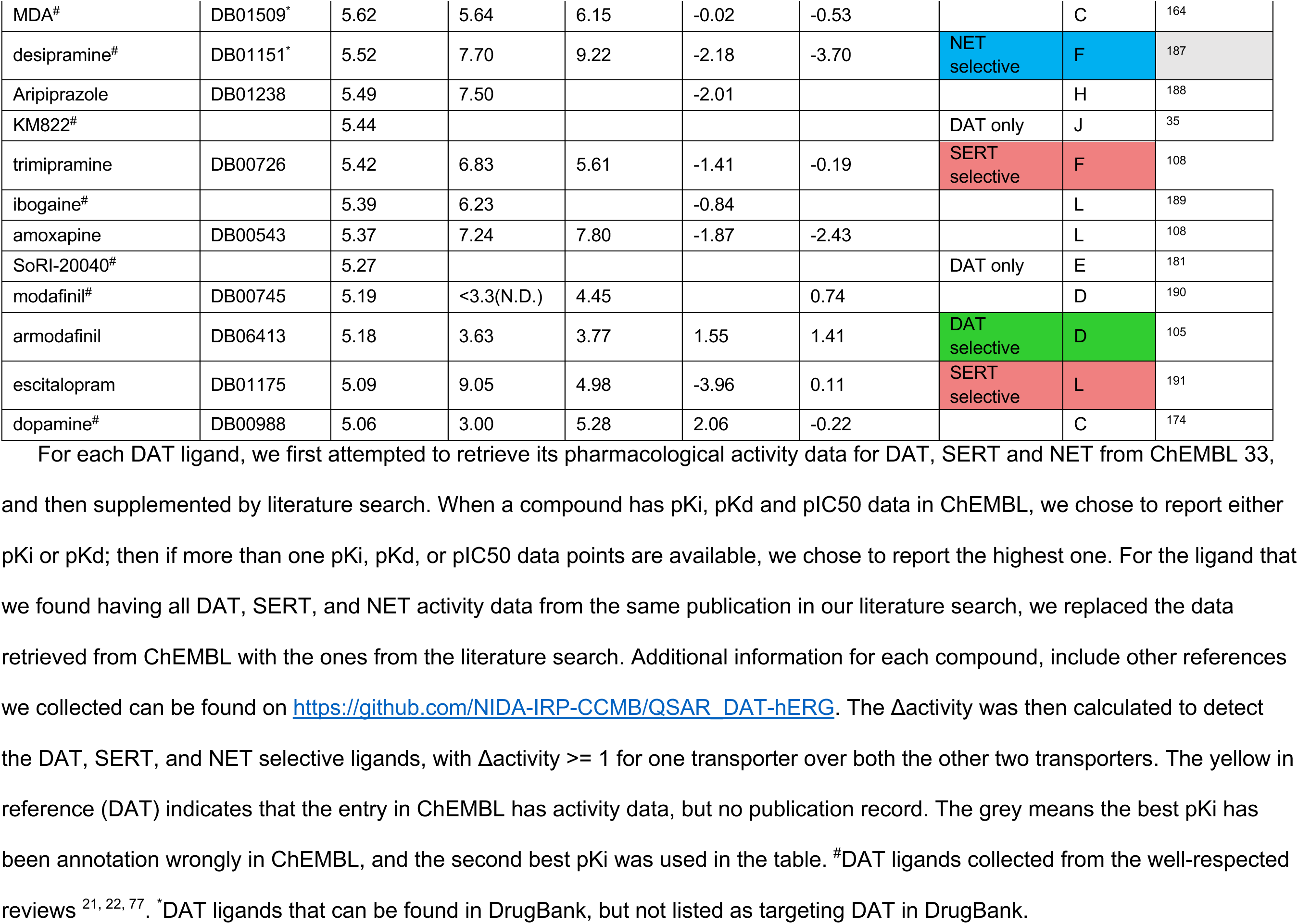
The pharmacological activities of some representative DAT ligands.

**Figure S1.**
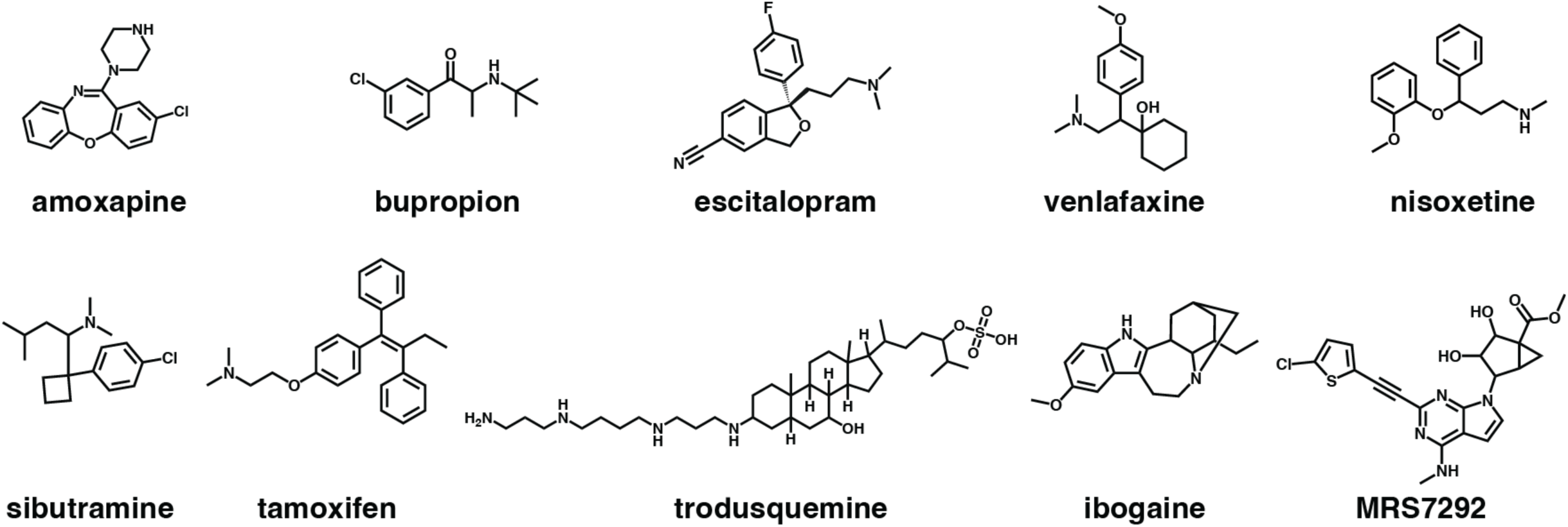
Single-member clusters.

## References

(1) Chen, N. H.; Reith, M. E.; Quick, M. W. Synaptic uptake and beyond: the sodium- and chloride-dependent neurotransmitter transporter family SLC6. Pflugers Arch 2004, 447 (5), 519–531. DOI: 10.1007/s00424-003-1064-5 From NLM Medline.

(2) Beuming, T.; Shi, L.; Javitch, J. A.; Weinstein, H. A comprehensive structure-based alignment of prokaryotic and eukaryotic neurotransmitter/Na+ symporters (NSS) aids in the use of the LeuT structure to probe NSS structure and function. Mol Pharmacol 2006, 70 (5), 1630–1642. DOI: 10.1124/mol.106.026120.

(3) Aggarwal, S.; Mortensen, O. V. Overview of Monoamine Transporters. Curr Protoc Pharmacol 2017, 79, 12 16 11-12 16 17. DOI: 10.1002/cpph.32 From NLM Medline.

(4) Juarez Olguin, H.; Calderon Guzman, D.; Hernandez Garcia, E.; Barragan Mejia, G. The Role of Dopamine and Its Dysfunction as a Consequence of Oxidative Stress. Oxid Med Cell Longev 2016, 2016, 9730467. DOI: 10.1155/2016/9730467 From NLM Medline.

(5) Stolzenberg, S.; Quick, M.; Zhao, C.; Gotfryd, K.; Khelashvili, G.; Gether, U.; Loland, C. J.; Javitch, J. A.; Noskov, S.; Weinstein, H.; Shi, L. Mechanism of the Association between Na+ Binding and Conformations at the Intracellular Gate in Neurotransmitter:Sodium Symporters. J Biol Chem 2015, 290 (22), 13992–14003. DOI: 10.1074/jbc.M114.625343.

(6) Borre, L.; Andreassen, T. F.; Shi, L.; Weinstein, H.; Gether, U. The second sodium site in the dopamine transporter controls cation permeation and is regulated by chloride. J Biol Chem 2014, 289 (37), 25764–25773. DOI: 10.1074/jbc.M114.574269.

(7) Cook, E. H., Jr.; Stein, M. A.; Krasowski, M. D.; Cox, N. J.; Olkon, D. M.; Kieffer, J. E.; Leventhal, B. L. Association of attention-deficit disorder and the dopamine transporter gene. Am J Hum Genet 1995, 56 (4), 993–998. From NLM Medline.

(8) Waldman, I. D.; Rowe, D. C.; Abramowitz, A.; Kozel, S. T.; Mohr, J. H.; Sherman, S. L.; Cleveland, H. H.; Sanders, M. L.; Gard, J. M.; Stever, C. Association and linkage of the dopamine transporter gene and attention-deficit hyperactivity disorder in children: heterogeneity owing to diagnostic subtype and severity. Am J Hum Genet 1998, 63 (6), 1767–1776. DOI: 10.1086/302132 From NLM Medline.

(9) Greenwood, T. A.; Schork, N. J.; Eskin, E.; Kelsoe, J. R. Identification of additional variants within the human dopamine transporter gene provides further evidence for an association with bipolar disorder in two independent samples. Mol Psychiatry 2006, 11 (2), 125–133, 115. DOI: 10.1038/sj.mp.4001764 From NLM Medline.

(10) Ashok, A. H.; Marques, T. R.; Jauhar, S.; Nour, M. M.; Goodwin, G. M.; Young, A. H.; Howes, O. D. The dopamine hypothesis of bipolar affective disorder: the state of the art and implications for treatment. Mol Psychiatry 2017, 22 (5), 666–679. DOI: 10.1038/mp.2017.16 From NLM Medline.

(11) Tamura, T.; Sugihara, G.; Okita, K.; Mukai, Y.; Matsuda, H.; Shiwaku, H.; Takagi, S.; Daisaki, H.; Tateishi, U.; Takahashi, H. Dopamine dysfunction in depression: application of texture analysis to dopamine transporter single-photon emission computed tomography imaging. Transl Psychiatry 2022, 12 (1), 309. DOI: 10.1038/s41398-022-02080-z From NLM Medline.

(12) McLellan, A. T. Substance Misuse and Substance use Disorders: Why do they Matter in Healthcare? Trans Am Clin Climatol Assoc 2017, 128, 112–130. From NLM Medline.

(13) Schwartz, E. K. C.; Wolkowicz, N. R.; De Aquino, J. P.; MacLean, R. R.; Sofuoglu, M. Cocaine Use Disorder (CUD): Current Clinical Perspectives. Subst Abuse Rehabil 2022, 13, 25–46. DOI: 10.2147/SAR.S337338 From NLM PubMed-not-MEDLINE.

(14) Abuse, N. I. o. D. Drug Overdose Death Rates. 2023. https://nida.nih.gov/research-topics/trends-statistics/overdose-death-rates (accessed December 16, 2023).

(15) Giros, B.; Jaber, M.; Jones, S. R.; Wightman, R. M.; Caron, M. G. Hyperlocomotion and indifference to cocaine and amphetamine in mice lacking the dopamine transporter. Nature 1996, 379 (6566), 606–612. DOI: 10.1038/379606a0 From NLM Medline.

(16) Chen, R.; Tilley, M. R.; Wei, H.; Zhou, F.; Zhou, F. M.; Ching, S.; Quan, N.; Stephens, R. L.; Hill, E. R.; Nottoli, T.;, et al. Abolished cocaine reward in mice with a cocaine-insensitive dopamine transporter. Proc Natl Acad Sci U S A 2006, 103 (24), 9333–9338. DOI: 10.1073/pnas.0600905103 From NLM Medline.

(17) Newman, A. H.; Kulkarni, S. Probes for the dopamine transporter: new leads toward a cocaine-abuse therapeutic--A focus on analogues of benztropine and rimcazole. Med Res Rev 2002, 22 (5), 429–464. DOI: 10.1002/med.10014 From NLM Medline.

(18) Newman, A. H.; Katz, J. L. Atypical Dopamine Uptake Inhibitors that Provide Clues About Cocaine’s Mechanism at the Dopamine Transporter. Top Med Chem Ser 2009, 4, 95–129. DOI: 10.1007/7355_2008_027.

(19) Loland, C. J.; Desai, R. I.; Zou, M. F.; Cao, J.; Grundt, P.; Gerstbrein, K.; Sitte, H. H.; Newman, A. H.; Katz, J. L.; Gether, U. Relationship between conformational changes in the dopamine transporter and cocaine-like subjective effects of uptake inhibitors. Mol Pharmacol 2008, 73 (3), 813–823. DOI: 10.1124/mol.107.039800 From NLM Medline.

(20) Reith, M. E.; Blough, B. E.; Hong, W. C.; Jones, K. T.; Schmitt, K. C.; Baumann, M. H.; Partilla, J. S.; Rothman, R. B.; Katz, J. L. Behavioral, biological, and chemical perspectives on atypical agents targeting the dopamine transporter. Drug Alcohol Depend 2015, 147, 1–19. DOI: 10.1016/j.drugalcdep.2014.12.005 From NLM Medline.

(21) Schmitt, K. C.; Reith, M. E. The atypical stimulant and nootropic modafinil interacts with the dopamine transporter in a different manner than classical cocaine-like inhibitors. PLoS One 2011, 6 (10), e25790. DOI: 10.1371/journal.pone.0025790 From NLM Medline.

(22) Nepal, B.; Das, S.; Reith, M. E.; Kortagere, S. Overview of the structure and function of the dopamine transporter and its protein interactions. Front Physiol 2023, 14, 1150355. DOI: 10.3389/fphys.2023.1150355 From NLM PubMed-not-MEDLINE.

(23) Abramyan, A. M.; Stolzenberg, S.; Li, Z.; Loland, C. J.; Noe, F.; Shi, L. The Isomeric Preference of an Atypical Dopamine Transporter Inhibitor Contributes to Its Selection of the Transporter Conformation. ACS Chem Neurosci 2017, 8 (8), 1735–1746. DOI: 10.1021/acschemneuro.7b00094 From NLM Medline.

(24) Zou, M. F.; Cao, J.; Abramyan, A. M.; Kopajtic, T.; Zanettini, C.; Guthrie, D. A.; Rais, R.; Slusher, B. S.; Shi, L.; Loland, C. J.; Newman, A. H. Structure-Activity Relationship Studies on a Series of 3alpha-[Bis(4-fluorophenyl)methoxy]tropanes and 3alpha-[Bis(4-fluorophenyl)methylamino]tropanes As Novel Atypical Dopamine Transporter (DAT) Inhibitors for the Treatment of Cocaine Use Disorders. J Med Chem 2017, 60 (24), 10172–10187. DOI: 10.1021/acs.jmedchem.7b01454 From NLM Medline.

(25) Rothman, R. B.; Baumann, M. H.; Prisinzano, T. E.; Newman, A. H. Dopamine transport inhibitors based on GBR12909 and benztropine as potential medications to treat cocaine addiction. Biochem Pharmacol 2008, 75 (1), 2–16. DOI: 10.1016/j.bcp.2007.08.007 From NLM Medline.

(26) Zhou, J.; He, R.; Johnson, K. M.; Ye, Y.; Kozikowski, A. P. Piperidine-based nocaine/modafinil hybrid ligands as highly potent monoamine transporter inhibitors: efficient drug discovery by rational lead hybridization. J Med Chem 2004, 47 (24), 5821–5824. DOI: 10.1021/jm040117o From NLM Medline.

(27) Loland, C. J.; Mereu, M.; Okunola, O. M.; Cao, J.; Prisinzano, T. E.; Mazier, S.; Kopajtic, T.; Shi, L.; Katz, J. L.; Tanda, G.; Newman, A. H. R-modafinil (armodafinil): a unique dopamine uptake inhibitor and potential medication for psychostimulant abuse. Biol Psychiatry 2012, 72 (5), 405–413. DOI: 10.1016/j.biopsych.2012.03.022 From NLM Medline.

(28) Okunola-Bakare, O. M.; Cao, J.; Kopajtic, T.; Katz, J. L.; Loland, C. J.; Shi, L.; Newman, A. H. Elucidation of structural elements for selectivity across monoamine transporters: novel 2-[(diphenylmethyl)sulfinyl]acetamide (modafinil) analogues. J Med Chem 2014, 57 (3), 1000–1013. DOI: 10.1021/jm401754x.

(29) Newman, A. H.; Ku, T.; Jordan, C. J.; Bonifazi, A.; Xi, Z. X. New Drugs, Old Targets: Tweaking the Dopamine System to Treat Psychostimulant Use Disorders. Annu Rev Pharmacol Toxicol 2021, 61 (1), 609–628. DOI: 10.1146/annurev-pharmtox-030220-124205 (acccessed 2023/12/16). From NLM Medline.

(30) Pratuangdejkul, J.; Schneider, B.; Launay, J. M.; Kellermann, O.; Manivet, P. Computational approaches for the study of serotonin and its membrane transporter SERT: implications for drug design in neurological sciences. Curr Med Chem 2008, 15 (30), 3214–3227. DOI: 10.2174/092986708786848523 From NLM Medline.

(31) Nolan, T. L.; Geffert, L. M.; Kolber, B. J.; Madura, J. D.; Surratt, C. K. Discovery of novel-scaffold monoamine transporter ligands via in silico screening with the S1 pocket of the serotonin transporter. ACS Chem Neurosci 2014, 5 (9), 784–792. DOI: 10.1021/cn500133b From NLM Medline.

(32) Manepalli, S.; Geffert, L. M.; Surratt, C. K.; Madura, J. D. Discovery of novel selective serotonin reuptake inhibitors through development of a protein-based pharmacophore. J Chem Inf Model 2011, 51 (9), 2417–2426. DOI: 10.1021/ci200280m From NLM Medline.

(33) Simmons, K. J.; Gotfryd, K.; Billesbolle, C. B.; Loland, C. J.; Gether, U.; Fishwick, C. W.; Johnson, A. P. A virtual high-throughput screening approach to the discovery of novel inhibitors of the bacterial leucine transporter, LeuT. Mol Membr Biol 2013, 30 (2), 184–194. DOI: 10.3109/09687688.2012.710341 From NLM Medline.

(34) Schlessinger, A.; Geier, E.; Fan, H.; Irwin, J. J.; Shoichet, B. K.; Giacomini, K. M.; Sali, A. Structure-based discovery of prescription drugs that interact with the norepinephrine transporter, NET. Proc Natl Acad Sci U S A 2011, 108 (38), 15810–15815. DOI: 10.1073/pnas.1106030108 From NLM Medline.

(35) Aggarwal, S.; Liu, X.; Rice, C.; Menell, P.; Clark, P. J.; Paparoidamis, N.; Xiao, Y. C.; Salvino, J. M.; Fontana, A. C. K.; Espana, R. A.;, et al. Identification of a Novel Allosteric Modulator of the Human Dopamine Transporter. ACS Chem Neurosci 2019, 10 (8), 3718–3730. DOI: 10.1021/acschemneuro.9b00262 From NLM Medline.

(36) Indarte, M.; Madura, J. D.; Surratt, C. K. Dopamine transporter comparative molecular modeling and binding site prediction using the LeuT(Aa) leucine transporter as a template. Proteins 2008, 70 (3), 1033–1046. DOI: 10.1002/prot.21598 From NLM Medline.

(37) Vemula, D.; Jayasurya, P.; Sushmitha, V.; Kumar, Y. N.; Bhandari, V. CADD, AI and ML in drug discovery: A comprehensive review. Eur J Pharm Sci 2023, 181, 106324. DOI: 10.1016/j.ejps.2022.106324 From NLM Medline.

(38) Voet, A.; Banwell, E. F.; Sahu, K. K.; Heddle, J. G.; Zhang, K. Y. Protein interface pharmacophore mapping tools for small molecule protein: protein interaction inhibitor discovery. Curr Top Med Chem 2013, 13 (9), 989–1001. DOI: 10.2174/1568026611313090003 From NLM Medline.

(39) Khedkar, S. A.; Malde, A. K.; Coutinho, E. C.; Srivastava, S. Pharmacophore modeling in drug discovery and development: an overview. Med Chem 2007, 3 (2), 187–197. DOI: 10.2174/157340607780059521 From NLM Medline.

(40) Christensen, H. S.; Boye, S. V.; Thinggaard, J.; Sinning, S.; Wiborg, O.; Schiott, B.; Bols, M. QSAR studies and pharmacophore identification for arylsubstituted cycloalkenecarboxylic acid methyl esters with affinity for the human dopamine transporter. Bioorg Med Chem 2007, 15 (15), 5262–5274. DOI: 10.1016/j.bmc.2007.05.015 From NLM Medline.

(41) Hansch, C.; Maloney, P. P.; Fujita, T. Correlation of Biological Activity of Phenoxyacetic Acids with Hammett Substituent Constants and Partition Coefficients. Nature 1962, 194 (4824), 178-&. DOI: DOI 10.1038/194178b0.

(42) Verma, R. P.; Hansch, C. Camptothecins: a SAR/QSAR study. Chem Rev 2009, 109 (1), 213–235. DOI: 10.1021/cr0780210 From NLM Medline.

(43) Gao, K.; Chen, D.; Robison, A. J.; Wei, G. W. Proteome-Informed Machine Learning Studies of Cocaine Addiction. J Phys Chem Lett 2021, 12 (45), 11122–11134. DOI: 10.1021/acs.jpclett.1c03133 From NLM Medline.

(44) Danishuddin; Khan, A. U. Descriptors and their selection methods in QSAR analysis: paradigm for drug design. Drug Discov Today 2016, 21 (8), 1291–1302. DOI: 10.1016/j.drudis.2016.06.013 From NLM Medline.

(45) Mao, J.; Akhtar, J.; Zhang, X.; Sun, L.; Guan, S.; Li, X.; Chen, G.; Liu, J.; Jeon, H. N.; Kim, M. S.;, et al. Comprehensive strategies of machine-learning-based quantitative structure-activity relationship models. iScience 2021, 24 (9), 103052. DOI: 10.1016/j.isci.2021.103052 From NLM PubMed-not-MEDLINE.

(46) Hong, H.; Xie, Q.; Ge, W.; Qian, F.; Fang, H.; Shi, L.; Su, Z.; Perkins, R.; Tong, W. Mold(2), molecular descriptors from 2D structures for chemoinformatics and toxicoinformatics. J Chem Inf Model 2008, 48 (7), 1337–1344. DOI: 10.1021/ci800038f From NLM Medline.

(47) Mauri, A.; Consonni, V.; Pavan, M.; Todeschini, R. Dragon software: An easy approach to molecular descriptor calculations. Match-Commun Math Co 2006, 56 (2), 237–248, Article. Scopus.

(48) Cho, Y. S.; No, K. T.; Cho, K. H. yaInChI: modified InChI string scheme for line notation of chemical structures. SAR QSAR Environ Res 2012, 23 (3-4), 237–255. DOI: 10.1080/1062936X.2012.657677 From NLM Medline.

(49) Choi, S. Y.; Shin, J. H.; Ryu, C. K.; Nam, K. Y.; No, K. T.; Park Choo, H. Y. The development of 3D-QSAR study and recursive partitioning of heterocyclic quinone derivatives with antifungal activity. Bioorg Med Chem 2006, 14 (5), 1608–1617. DOI: 10.1016/j.bmc.2005.10.010 From NLM Medline.

(50) Hayakawa, D.; Sawada, N.; Watanabe, Y.; Gouda, H. A molecular interaction field describing nonconventional intermolecular interactions and its application to protein-ligand interaction prediction. J Mol Graph Model 2020, 96, 107515. DOI: 10.1016/j.jmgm.2019.107515 From NLM Medline.

(51) Hoffman, B. T.; Kopajtic, T.; Katz, J. L.; Newman, A. H. 2D QSAR modeling and preliminary database searching for dopamine transporter inhibitors using genetic algorithm variable selection of Molconn Z descriptors. J Med Chem 2000, 43 (22), 4151–4159. DOI: 10.1021/jm990472s From NLM Medline.

(52) Kulkarni, S. S.; Newman, A. H.; Houlihan, W. J. Three-dimensional quantitative structure-activity relationships of mazindol analogues at the dopamine transporter. J Med Chem 2002, 45 (19), 4119–4127. DOI: 10.1021/jm0102093 From NLM Medline.

(53) Zhang, X.; Mao, J.; Wei, M.; Qi, Y.; Zhang, J. Z. H. HergSPred: Accurate Classification of hERG Blockers/Nonblockers with Machine-Learning Models. J Chem Inf Model 2022, 62 (8), 1830–1839. DOI: 10.1021/acs.jcim.2c00256 From NLM Medline.

(54) Soares, T. A.; Nunes-Alves, A.; Mazzolari, A.; Ruggiu, F.; Wei, G. W.; Merz, K. The (Re)-Evolution of Quantitative Structure-Activity Relationship (QSAR) Studies Propelled by the Surge of Machine Learning Methods. J Chem Inf Model 2022, 62 (22), 5317–5320. DOI: 10.1021/acs.jcim.2c01422 From NLM Medline.

(55) Alsenan, S. A.; Al-Turaiki, I. M.; Hafez, A. M. Feature Extraction Methods in Quantitative Structure–Activity Relationship Modeling: A Comparative Study. IEEE Access 2020, 8, 78737–78752. DOI: 10.1109/access.2020.2990375.

(56) Tropsha, A.; Isayev, O.; Varnek, A.; Schneider, G.; Cherkasov, A. Integrating QSAR modelling and deep learning in drug discovery: the emergence of deep QSAR. Nat Rev Drug Discov 2023. DOI: 10.1038/s41573-023-00832-0 From NLM Publisher.

(57) Yao, X. J.; Panaye, A.; Doucet, J. P.; Zhang, R. S.; Chen, H. F.; Liu, M. C.; Hu, Z. D.; Fan, B. T. Comparative study of QSAR/QSPR correlations using support vector machines, radial basis function neural networks, and multiple linear regression. J Chem Inf Comput Sci 2004, 44 (4), 1257–1266. DOI: 10.1021/ci049965i From NLM Medline.

(58) Ma, J.; Sheridan, R. P.; Liaw, A.; Dahl, G. E.; Svetnik, V. Deep neural nets as a method for quantitative structure-activity relationships. J Chem Inf Model 2015, 55 (2), 263–274. DOI: 10.1021/ci500747n From NLM Medline.

(59) Sheridan, R. P.; Wang, W. M.; Liaw, A.; Ma, J.; Gifford, E. M. Extreme Gradient Boosting as a Method for Quantitative Structure-Activity Relationships. J Chem Inf Model 2016, 56 (12), 2353–2360. DOI: 10.1021/acs.jcim.6b00591 From NLM Medline.

(60) Svetnik, V.; Liaw, A.; Tong, C.; Culberson, J. C.; Sheridan, R. P.; Feuston, B. P. Random forest: a classification and regression tool for compound classification and QSAR modeling. J Chem Inf Comput Sci 2003, 43 (6), 1947–1958. DOI: 10.1021/ci034160g From NLM PubMed-not-MEDLINE.

(61) Gaulton, A.; Bellis, L. J.; Bento, A. P.; Chambers, J.; Davies, M.; Hersey, A.; Light, Y.; McGlinchey, S.; Michalovich, D.; Al-Lazikani, B.; Overington, J. P. ChEMBL: a large-scale bioactivity database for drug discovery. Nucleic Acids Res 2012, 40 (Database issue), D1100–1107. DOI: 10.1093/nar/gkr777 From NLM Medline.

(62) Wishart, D. S.; Feunang, Y. D.; Guo, A. C.; Lo, E. J.; Marcu, A.; Grant, J. R.; Sajed, T.; Johnson, D.; Li, C.; Sayeeda, Z.;, et al. DrugBank 5.0: a major update to the DrugBank database for 2018. Nucleic Acids Res 2018, 46 (D1), D1074–D1082. DOI: 10.1093/nar/gkx1037 From NLM Medline.

(63) Gilson, M. K.; Liu, T.; Baitaluk, M.; Nicola, G.; Hwang, L.; Chong, J. BindingDB in 2015: A public database for medicinal chemistry, computational chemistry and systems pharmacology. Nucleic Acids Res 2016, 44 (D1), D1045-1053. DOI: 10.1093/nar/gkv1072 From NLM Medline.

(64) Liu, T.; Lin, Y.; Wen, X.; Jorissen, R. N.; Gilson, M. K. BindingDB: a web-accessible database of experimentally determined protein-ligand binding affinities. Nucleic Acids Res 2007, 35 (Database issue), D198-201. DOI: 10.1093/nar/gkl999 From NLM Medline.

(65) Zhou, Y.; Zhang, Y.; Zhao, D.; Yu, X.; Shen, X.; Zhou, Y.; Wang, S.; Qiu, Y.; Chen, Y.; Zhu, F. TTD: Therapeutic Target Database describing target druggability information. Nucleic Acids Res 2024, 52 (D1), D1465–D1477. DOI: 10.1093/nar/gkad751 From NLM Medline.

(66) Besnard, J.; Ruda, G. F.; Setola, V.; Abecassis, K.; Rodriguiz, R. M.; Huang, X. P.; Norval, S.; Sassano, M. F.; Shin, A. I.; Webster, L. A.;, et al. Automated design of ligands to polypharmacological profiles. Nature 2012, 492 (7428), 215–220. DOI: 10.1038/nature11691 From NLM Medline.

(67) Kim, S.; Chen, J.; Cheng, T.; Gindulyte, A.; He, J.; He, S.; Li, Q.; Shoemaker, B. A.; Thiessen, P. A.; Yu, B.;, et al. PubChem 2023 update. Nucleic Acids Res 2023, 51 (D1), D1373-D1380. DOI: 10.1093/nar/gkac956 From NLM Medline.

(68) Lee, K. H.; Fant, A. D.; Guo, J.; Guan, A.; Jung, J.; Kudaibergenova, M.; Miranda, W. E.; Ku, T.; Cao, J.; Wacker, S.;, et al. Toward Reducing hERG Affinities for DAT Inhibitors with a Combined Machine Learning and Molecular Modeling Approach. J Chem Inf Model 2021, 61 (9), 4266–4279. DOI: 10.1021/acs.jcim.1c00856 From NLM Medline.

(69) Tuomisto, J.; Tuomisto, L.; Pazdernik, T. L. Conformationally rigid amphetamine analogs as inhibitors of monoamine uptake by brain synaptosomes. J Med Chem 1976, 19 (5), 725–727. DOI: 10.1021/jm00227a030 From NLM Medline.

(70) Yamashita, A.; Singh, S. K.; Kawate, T.; Jin, Y.; Gouaux, E. Crystal structure of a bacterial homologue of Na+/Cl--dependent neurotransmitter transporters. Nature 2005, 437 (7056), 215–223. DOI: 10.1038/nature03978 From NLM Medline.

(71) Koldso, H.; Christiansen, A. B.; Sinning, S.; Schiott, B. Comparative modeling of the human monoamine transporters: similarities in substrate binding. ACS Chem Neurosci 2013, 4 (2), 295–309. DOI: 10.1021/cn300148r From NLM Medline.

(72) Severinsen, K.; Koldso, H.; Thorup, K. A.; Schjoth-Eskesen, C.; Moller, P. T.; Wiborg, O.; Jensen, H. H.; Sinning, S.; Schiott, B. Binding of mazindol and analogs to the human serotonin and dopamine transporters. Mol Pharmacol 2014, 85 (2), 208–217. DOI: 10.1124/mol.113.088922 From NLM Medline.

(73) Shan, J.; Javitch, J. A.; Shi, L.; Weinstein, H. The substrate-driven transition to an inward-facing conformation in the functional mechanism of the dopamine transporter. PLoS One 2011, 6 (1), e16350. DOI: 10.1371/journal.pone.0016350 From NLM Medline.

(74) Cheng, M. H.; Bahar, I. Molecular Mechanism of Dopamine Transport by Human Dopamine Transporter. Structure 2015, 23 (11), 2171–2181. DOI: 10.1016/j.str.2015.09.001.

(75) Ravna, A. W.; Sylte, I.; Dahl, S. G. Structure and localisation of drug binding sites on neurotransmitter transporters. J Mol Model 2009, 15 (10), 1155–1164. DOI: 10.1007/s00894-009-0478-1 From NLM Medline.

(76) Penmatsa, A.; Wang, K. H.; Gouaux, E. X-ray structure of dopamine transporter elucidates antidepressant mechanism. Nature 2013, 503 (7474), 85–90. DOI: 10.1038/nature12533 From NLM Medline.

(77) Schmitt, K. C.; Rothman, R. B.; Reith, M. E. Nonclassical pharmacology of the dopamine transporter: atypical inhibitors, allosteric modulators, and partial substrates. J Pharmacol Exp Ther 2013, 346 (1), 2–10. DOI: 10.1124/jpet.111.191056 From NLM Medline.

(78) Gatley, S. J.; Volkow, N. D.; Chen, R.; Fowler, J. S.; Carroll, F. I.; Kuhar, M. J. Displacement of RTI-55 from the dopamine transporter by cocaine. Eur J Pharmacol 1996, 296 (2), 145–151. DOI: 10.1016/0014-2999(95)00698-2 From NLM Medline.

(79) Coffey, L. L.; Reith, M. E. [3H]WIN 35,428 binding to the dopamine uptake carrier. I. Effect of tonicity and buffer composition. J Neurosci Methods 1994, 51 (1), 23–30. DOI: 10.1016/0165-0270(94)90022-1 From NLM Medline.

(80) Dahal, R. A.; Pramod, A. B.; Sharma, B.; Krout, D.; Foster, J. D.; Cha, J. H.; Cao, J.; Newman, A. H.; Lever, J. R.; Vaughan, R. A.; Henry, L. K. Computational and biochemical docking of the irreversible cocaine analog RTI 82 directly demonstrates ligand positioning in the dopamine transporter central substrate-binding site. J Biol Chem 2014, 289 (43), 29712–29727. DOI: 10.1074/jbc.M114.571521 From NLM Medline.

(81) Parnas, M. L.; Gaffaney, J. D.; Zou, M. F.; Lever, J. R.; Newman, A. H.; Vaughan, R. A. Labeling of dopamine transporter transmembrane domain 1 with the tropane ligand N-[4-(4-azido-3-[125I]iodophenyl)butyl]-2beta-carbomethoxy-3beta-(4-chlorophenyl)tropane implicates proximity of cocaine and substrate active sites. Mol Pharmacol 2008, 73 (4), 1141–1150. DOI: 10.1124/mol.107.043679 From NLM Medline.

(82) Booij, J.; Andringa, G.; Rijks, L. J.; Vermeulen, R. J.; De Bruin, K.; Boer, G. J.; Janssen, A. G.; Van Royen, E. A. [123I]FP-CIT binds to the dopamine transporter as assessed by biodistribution studies in rats and SPECT studies in MPTP-lesioned monkeys. Synapse 1997, 27 (3), 183–190. DOI: 10.1002/(SICI)1098-2396(199711)27:3<183::AID-SYN4>3.0.CO;2-9 From NLM Medline.

(83) Madras, B. K.; Meltzer, P. C.; Liang, A. Y.; Elmaleh, D. R.; Babich, J.; Fischman, A. J. Altropane, a SPECT or PET imaging probe for dopamine neurons: I. Dopamine transporter binding in primate brain. Synapse 1998, 29 (2), 93–104. DOI: 10.1002/(SICI)1098-2396(199806)29:2<93::AID-SYN1>3.0.CO;2-5 From NLM Medline.

(84) Kimmel, H. L.; Carroll, F. I.; Kuhar, M. J. Locomotor stimulant effects of novel phenyltropanes in the mouse. Drug Alcohol Depend 2001, 65 (1), 25–36. DOI: 10.1016/s0376-8716(01)00144-2 From NLM Medline.

(85) Lehr, T.; Staab, A.; Tillmann, C.; Nielsen, E. O.; Trommeshauser, D.; Schaefer, H. G.; Kloft, C. Contribution of the active metabolite M1 to the pharmacological activity of tesofensine in vivo: a pharmacokinetic-pharmacodynamic modelling approach. Br J Pharmacol 2008, 153 (1), 164–174. DOI: 10.1038/sj.bjp.0707539 From NLM Medline.

(86) Elia, J.; Ambrosini, P. J.; Rapoport, J. L. Treatment of attention-deficit-hyperactivity disorder. N Engl J Med 1999, 340 (10), 780–788. DOI: 10.1056/NEJM199903113401007 From NLM Medline.

(87) Kutcher, S.; Aman, M.; Brooks, S. J.; Buitelaar, J.; van Daalen, E.; Fegert, J.; Findling, R. L.; Fisman, S.; Greenhill, L. L.; Huss, M.;, et al. International consensus statement on attention-deficit/hyperactivity disorder (ADHD) and disruptive behaviour disorders (DBDs): clinical implications and treatment practice suggestions. Eur Neuropsychopharmacol 2004, 14 (1), 11–28. DOI: 10.1016/s0924-977x(03)00045-2 From NLM Medline.

(88) Barch, D. M.; Carter, C. S. Amphetamine improves cognitive function in medicated individuals with schizophrenia and in healthy volunteers. Schizophr Res 2005, 77 (1), 43–58. DOI: 10.1016/j.schres.2004.12.019 From NLM Medline.

(89) Berman, S. M.; Kuczenski, R.; McCracken, J. T.; London, E. D. Potential adverse effects of amphetamine treatment on brain and behavior: a review. Mol Psychiatry 2009, 14 (2), 123–142. DOI: 10.1038/mp.2008.90 From NLM Medline.

(90) Northrop, N. A.; Yamamoto, B. K. Methamphetamine effects on blood-brain barrier structure and function. Front Neurosci 2015, 9, 69. DOI: 10.3389/fnins.2015.00069 From NLM PubMed-not-MEDLINE.

(91) McCann, U. D.; Wong, D. F.; Yokoi, F.; Villemagne, V.; Dannals, R. F.; Ricaurte, G. A. Reduced striatal dopamine transporter density in abstinent methamphetamine and methcathinone users: evidence from positron emission tomography studies with [11C]WIN-35,428. J Neurosci 1998, 18 (20), 8417–8422. DOI: 10.1523/JNEUROSCI.18-20-08417.1998 From NLM Medline.

(92) Paulus, M. P.; Stewart, J. L. Neurobiology, Clinical Presentation, and Treatment of Methamphetamine Use Disorder: A Review. JAMA Psychiatry 2020, 77 (9), 959–966. DOI: 10.1001/jamapsychiatry.2020.0246 From NLM Medline.

(93) Martin, D., Le, J. K.. Amphetamine [Updated 2023 Jul 31]. In: StatPearls [Internet]. 2023. https://www.ncbi.nlm.nih.gov/books/NBK556103/ (accessed.

(94) Erdo, S. L.; Kiss, B.; Rosdy, B. Inhibition of dopamine uptake by a new psychostimulant mesocarb (Sydnocarb). Pol J Pharmacol Pharm 1981, 33 (2), 141–147. From NLM Medline.

(95) Aggarwal, S.; Cheng, M. H.; Salvino, J. M.; Bahar, I.; Mortensen, O. V. Functional Characterization of the Dopaminergic Psychostimulant Sydnocarb as an Allosteric Modulator of the Human Dopamine Transporter. Biomedicines 2021, 9 (6). DOI: 10.3390/biomedicines9060634 From NLM PubMed-not-MEDLINE.

(96) Kalant, H. The pharmacology and toxicology of "ecstasy" (MDMA) and related drugs. CMAJ 2001, 165 (7), 917–928. From NLM Medline.

(97) Steinkellner, T.; Freissmuth, M.; Sitte, H. H.; Montgomery, T. The ugly side of amphetamines: short- and long-term toxicity of 3,4-methylenedioxymethamphetamine (MDMA, ‘Ecstasy’), methamphetamine and D-amphetamine. Biol Chem 2011, 392 (1-2), 103–115. DOI: 10.1515/BC.2011.016 From NLM Medline.

(98) Hagino, Y.; Takamatsu, Y.; Yamamoto, H.; Iwamura, T.; Murphy, D. L.; Uhl, G. R.; Sora, I.; Ikeda, K. Effects of MDMA on Extracellular Dopamine and Serotonin Levels in Mice Lacking Dopamine and/or Serotonin Transporters. Curr Neuropharmacol 2011, 9 (1), 91–95. DOI: 10.2174/157015911795017254 From NLM PubMed-not-MEDLINE.

(99) Rothman, R. B.; Partilla, J. S.; Baumann, M. H.; Lightfoot-Siordia, C.; Blough, B. E. Studies of the biogenic amine transporters. 14. Identification of low-efficacy "partial" substrates for the biogenic amine transporters. J Pharmacol Exp Ther 2012, 341 (1), 251-262. DOI: 10.1124/jpet.111.188946 From NLM Medline.

(100) Rothman, R. B.; Baumann, M. H. Therapeutic potential of monoamine transporter substrates. Curr Top Med Chem 2006, 6 (17), 1845–1859. DOI: 10.2174/156802606778249766 From NLM Medline.

(101) Lopez-Arnau, R.; Duart-Castells, L.; Aster, B.; Camarasa, J.; Escubedo, E.; Pubill, D. Effects of MDPV on dopamine transporter regulation in male rats. Comparison with cocaine. Psychopharmacology (Berl) 2019, 236 (3), 925–938. DOI: 10.1007/s00213-018-5052-z From NLM Medline.

(102) Baumann, M. H.; Partilla, J. S.; Lehner, K. R.; Thorndike, E. B.; Hoffman, A. F.; Holy, M.; Rothman, R. B.; Goldberg, S. R.; Lupica, C. R.; Sitte, H. H.;, et al. Powerful cocaine-like actions of 3,4-methylenedioxypyrovalerone (MDPV), a principal constituent of psychoactive ‘bath salts’ products. Neuropsychopharmacology 2013, 38 (4), 552–562. DOI: 10.1038/npp.2012.204 From NLM Medline.

(103) Baumann, M. H.; Wang, X.; Rothman, R. B. 3,4-Methylenedioxymethamphetamine (MDMA) neurotoxicity in rats: a reappraisal of past and present findings. Psychopharmacology (Berl) 2007, 189 (4), 407–424. DOI: 10.1007/s00213-006-0322-6 From NLM Medline.

(104) Volkow, N. D.; Fowler, J. S.; Logan, J.; Alexoff, D.; Zhu, W.; Telang, F.; Wang, G. J.; Jayne, M.; Hooker, J. M.; Wong, C.;, et al. Effects of modafinil on dopamine and dopamine transporters in the male human brain: clinical implications. JAMA 2009, 301 (11), 1148–1154. DOI: 10.1001/jama.2009.351 From NLM Medline.

(105) Shanmugasundaram, B.; Aher, Y. D.; Aradska, J.; Ilic, M.; Daba Feyissa, D.; Kalaba, P.; Aher, N. Y.; Dragacevic, V.; Saber Marouf, B.; Langer, T.;, et al. R-Modafinil exerts weak effects on spatial memory acquisition and dentate gyrus synaptic plasticity. PLoS One 2017, 12 (6), e0179675. DOI: 10.1371/journal.pone.0179675 From NLM Medline.

(106) Rotolo, R. A.; Dragacevic, V.; Kalaba, P.; Urban, E.; Zehl, M.; Roller, A.; Wackerlig, J.; Langer, T.; Pistis, M.; De Luca, M. A.;, et al. The Novel Atypical Dopamine Uptake Inhibitor (S)- CE-123 Partially Reverses the Effort-Related Effects of the Dopamine Depleting Agent Tetrabenazine and Increases Progressive Ratio Responding. Front Pharmacol 2019, 10, 682. DOI: 10.3389/fphar.2019.00682 From NLM PubMed-not-MEDLINE.

(107) Oleson, E. B.; Ferris, M. J.; Espana, R. A.; Harp, J.; Jones, S. R. Effects of the histamine H(1) receptor antagonist and benztropine analog diphenylpyraline on dopamine uptake, locomotion and reward. Eur J Pharmacol 2012, 683 (1-3), 161–165. DOI: 10.1016/j.ejphar.2012.03.003 From NLM Medline.

(108) Tatsumi, M.; Groshan, K.; Blakely, R. D.; Richelson, E. Pharmacological profile of antidepressants and related compounds at human monoamine transporters. Eur J Pharmacol 1997, 340 (2-3), 249–258. DOI: 10.1016/s0014-2999(97)01393-9 From NLM Medline.

(109) Newman, A. H.; Agoston, G. E. Novel benztropine [3a-(diphenylmethoxy)tropane] analogs as probes for the dopamine transporter. Curr Med Chem 1998, 5 (4), 305–319. From NLM Medline.

(110) Kopajtic, T. A.; Liu, Y.; Surratt, C. K.; Donovan, D. M.; Newman, A. H.; Katz, J. L. Dopamine transporter-dependent and -independent striatal binding of the benztropine analog JHW 007, a cocaine antagonist with low abuse liability. J Pharmacol Exp Ther 2010, 335 (3), 703–714. DOI: 10.1124/jpet.110.171629 From NLM Medline.

(111) Moerke, M. J.; Ananthan, S.; Banks, M. L.; Eltit, J. M.; Freitas, K. C.; Johnson, A. R.; Saini, S. K.; Steele, T. W. E.; Negus, S. S. Interactions between Cocaine and the Putative Allosteric Dopamine Transporter Ligand SRI-31142. J Pharmacol Exp Ther 2018, 367 (2), 222–233. DOI: 10.1124/jpet.118.250902 From NLM Medline.

(112) Rothman, R. B.; Dersch, C. M.; Carroll, F. I.; Ananthan, S. Studies of the biogenic amine transporters. VIII: identification of a novel partial inhibitor of dopamine uptake and dopamine transporter binding. Synapse 2002, 43 (4), 268–274. DOI: 10.1002/syn.10046 From NLM Medline.

(113) Pariser, J. J.; Partilla, J. S.; Dersch, C. M.; Ananthan, S.; Rothman, R. B. Studies of the biogenic amine transporters. 12. Identification of novel partial inhibitors of amphetamine-induced dopamine release. J Pharmacol Exp Ther 2008, 326 (1), 286–295. DOI: 10.1124/jpet.108.139675 From NLM Medline.

(114) Owens, M. J.; Morgan, W. N.; Plott, S. J.; Nemeroff, C. B. Neurotransmitter receptor and transporter binding profile of antidepressants and their metabolites. J Pharmacol Exp Ther 1997, 283 (3), 1305–1322. From NLM Medline.

(115) Goodnick, P. J.; Goldstein, B. J. Selective serotonin reuptake inhibitors in affective disorders--I. Basic pharmacology. J Psychopharmacol 1998, 12 (3 Suppl B), S5–20. DOI: 10.1177/0269881198012003021 From NLM Medline.

(116) Koblan, K. S.; Hopkins, S. C.; Sarma, K.; Jin, F.; Goldman, R.; Kollins, S. H.; Loebel, A. Dasotraline for the Treatment of Attention-Deficit/Hyperactivity Disorder: A Randomized, Double-Blind, Placebo-Controlled, Proof-of-Concept Trial in Adults. Neuropsychopharmacology 2015, 40 (12), 2745–2752. DOI: 10.1038/npp.2015.124 From NLM Medline.

(117) Lawler, C. P.; Prioleau, C.; Lewis, M. M.; Mak, C.; Jiang, D.; Schetz, J. A.; Gonzalez, A. M.; Sibley, D. R.; Mailman, R. B. Interactions of the novel antipsychotic aripiprazole (OPC-14597) with dopamine and serotonin receptor subtypes. Neuropsychopharmacology 1999, 20 (6), 612–627. DOI: 10.1016/S0893-133X(98)00099-2 From NLM Medline.

(118) Davies, M. A.; Sheffler, D. J.; Roth, B. L. Aripiprazole: a novel atypical antipsychotic drug with a uniquely robust pharmacology. CNS Drug Rev 2004, 10 (4), 317–336. DOI: 10.1111/j.1527-3458.2004.tb00030.x From NLM Medline.

(119) Duggal, H. S. Aripiprazole-induced improvement in tardive dyskinesia. Can J Psychiatry 2003, 48 (11), 771–772. DOI: 10.1177/070674370304801116 From NLM Medline.

(120) Carlier, P. R.; Lo, M. M.; Lo, P. C.; Richelson, E.; Tatsumi, M.; Reynolds, I. J.; Sharma, T. A. Synthesis of a potent wide-spectrum serotonin-, norepinephrine-, dopamine-reuptake inhibitor (SNDRI) and a species-selective dopamine-reuptake inhibitor based on the gamma-amino alcohol functional group. Bioorg Med Chem Lett 1998, 8 (5), 487–492. DOI: 10.1016/s0960-894x(98)00062-6 From NLM Medline.

(121) Karpa, K. D.; Cavanaugh, J. E.; Lakoski, J. M. Duloxetine pharmacology: profile of a dual monoamine modulator. CNS Drug Rev 2002, 8 (4), 361–376. DOI: 10.1111/j.1527-3458.2002.tb00234.x From NLM Medline.

(122) Cashman, J. R.; Ghirmai, S. Inhibition of serotonin and norepinephrine reuptake and inhibition of phosphodiesterase by multi-target inhibitors as potential agents for depression. Bioorg Med Chem 2009, 17 (19), 6890–6897. DOI: 10.1016/j.bmc.2009.08.025 From NLM Medline.

(123) Rothman, R. B.; Blough, B. E.; Woolverton, W. L.; Anderson, K. G.; Negus, S. S.; Mello, N. K.; Roth, B. L.; Baumann, M. H. Development of a rationally designed, low abuse potential, biogenic amine releaser that suppresses cocaine self-administration. J Pharmacol Exp Ther 2005, 313 (3), 1361–1369. DOI: 10.1124/jpet.104.082503 From NLM Medline.

(124) Hasenhuetl, P. S.; Bhat, S.; Freissmuth, M.; Sandtner, W. Functional Selectivity and Partial Efficacy at the Monoamine Transporters: A Unified Model of Allosteric Modulation and Amphetamine-Induced Substrate Release. Mol Pharmacol 2019, 95 (3), 303–312. DOI: 10.1124/mol.118.114793 From NLM Medline.

(125) Rothman, R. B.; Katsnelson, M.; Vu, N.; Partilla, J. S.; Dersch, C. M.; Blough, B. E.; Baumann, M. H. Interaction of the anorectic medication, phendimetrazine, and its metabolites with monoamine transporters in rat brain. Eur J Pharmacol 2002, 447 (1), 51–57. DOI: 10.1016/s0014-2999(02)01830-7 From NLM Medline.

(126) Fone, K. C.; Nutt, D. J. Stimulants: use and abuse in the treatment of attention deficit hyperactivity disorder. Curr Opin Pharmacol 2005, 5 (1), 87–93. DOI: 10.1016/j.coph.2004.10.001 From NLM Medline.

(127) Marco, P.; Maddalena, M.; Silvana, B.; Erika, G.; Maria, N. Attention Deficit Hyperactivity Disorder. In Comprehensive Pharmacology, 2022; pp 256–285.

(128) Raffel, D. M.; Chen, W. Binding of [3H]mazindol to cardiac norepinephrine transporters: kinetic and equilibrium studies. Naunyn Schmiedebergs Arch Pharmacol 2004, 370 (1), 9–16. DOI: 10.1007/s00210-004-0949-y From NLM Medline.

(129) Hiranita, T.; Soto, P. L.; Kohut, S. J.; Kopajtic, T.; Cao, J.; Newman, A. H.; Tanda, G.; Katz, J. L. Decreases in cocaine self-administration with dual inhibition of the dopamine transporter and sigma receptors. J Pharmacol Exp Ther 2011, 339 (2), 662–677. DOI: 10.1124/jpet.111.185025 From NLM Medline.

(130) Clark, A.; Tate, B.; Urban, B.; Schroeder, R.; Gennuso, S.; Ahmadzadeh, S.; McGregor, D.; Girma, B.; Shekoohi, S.; Kaye, A. D. Bupropion Mediated Effects on Depression, Attention Deficit Hyperactivity Disorder, and Smoking Cessation. Health Psychol Res 2023, 11, 81043. DOI: 10.52965/001c.81043 From NLM PubMed-not-MEDLINE.

(131) Mikelman, S. R.; Guptaroy, B.; Schmitt, K. C.; Jones, K. T.; Zhen, J.; Reith, M. E. A.; Gnegy, M. E. Tamoxifen Directly Interacts with the Dopamine Transporter. J Pharmacol Exp Ther 2018, 367 (1), 119–128. DOI: 10.1124/jpet.118.248179 From NLM Medline.

(132) Bulling, S.; Schicker, K.; Zhang, Y. W.; Steinkellner, T.; Stockner, T.; Gruber, C. W.; Boehm, S.; Freissmuth, M.; Rudnick, G.; Sitte, H. H.; Sandtner, W. The mechanistic basis for noncompetitive ibogaine inhibition of serotonin and dopamine transporters. J Biol Chem 2012, 287 (22), 18524–18534. DOI: 10.1074/jbc.M112.343681 From NLM Medline.

(133) Cheng, Y.; Prusoff, W. H. Relationship between the inhibition constant (K1) and the concentration of inhibitor which causes 50 per cent inhibition (I50) of an enzymatic reaction. Biochem Pharmacol 1973, 22 (23), 3099–3108. DOI: 10.1016/0006-2952(73)90196-2 From NLM Medline.

(134) Jumper, J.; Evans, R.; Pritzel, A.; Green, T.; Figurnov, M.; Ronneberger, O.; Tunyasuvunakool, K.; Bates, R.; Zidek, A.; Potapenko, A.;, et al. Highly accurate protein structure prediction with AlphaFold. Nature 2021, 596 (7873), 583–589. DOI: 10.1038/s41586-021-03819-2 From NLM Medline.

(135) AlQuraishi, M. Machine learning in protein structure prediction. Curr Opin Chem Biol 2021, 65, 1–8. DOI: 10.1016/j.cbpa.2021.04.005 From NLM Medline.

(136) Wenzel, J.; Matter, H.; Schmidt, F. Predictive Multitask Deep Neural Network Models for ADME-Tox Properties: Learning from Large Data Sets. J Chem Inf Model 2019, 59 (3), 1253–1268. DOI: 10.1021/acs.jcim.8b00785 From NLM Medline.

(137) Wu, K.; Wei, G. W. Quantitative Toxicity Prediction Using Topology Based Multitask Deep Neural Networks. J Chem Inf Model 2018, 58 (2), 520–531. DOI: 10.1021/acs.jcim.7b00558 From NLM Medline.

(138) Salas-Estrada, L.; Provasi, D.; Qiu, X.; Kaniskan, H. U.; Huang, X. P.; DiBerto, J. F.; Lamim Ribeiro, J. M.; Jin, J.; Roth, B. L.; Filizola, M. De Novo Design of kappa-Opioid Receptor Antagonists Using a Generative Deep-Learning Framework. J Chem Inf Model 2023, 63 (16), 5056–5065. DOI: 10.1021/acs.jcim.3c00651 From NLM Medline.

(139) Skinnider, M. A.; Wang, F.; Pasin, D.; Greiner, R.; Foster, L. J.; Dalsgaard, P. W.; Wishart, D. S. A deep generative model enables automated structure elucidation of novel psychoactive substances. Nat Mach Intell 2021, 3 (11), 973-+. DOI: 10.1038/s42256-021-00407-x.

(140) Atance, S. R.; Diez, J. V.; Engkvist, O.; Olsson, S.; Mercado, R. De Novo Drug Design Using Reinforcement Learning with Graph-Based Deep Generative Models. J Chem Inf Model 2022, 62 (20), 4863–4872. DOI: 10.1021/acs.jcim.2c00838 From NLM Medline.

(141) Kumar, R.; Sharma, A.; Alexiou, A.; Bilgrami, A. L.; Kamal, M. A.; Ashraf, G. M. DeePred-BBB: A Blood Brain Barrier Permeability Prediction Model With Improved Accuracy. Front Neurosci 2022, 16, 858126. DOI: 10.3389/fnins.2022.858126 From NLM PubMed-not-MEDLINE.

(142) Xu, Y. Deep Neural Networks for QSAR. Methods Mol Biol 2022, 2390, 233–260. DOI: 10.1007/978-1-0716-1787-8_10 From NLM Medline.

(143) Siramshetty, V. B.; Nguyen, D. T.; Martinez, N. J.; Southall, N. T.; Simeonov, A.; Zakharov, A. V. Critical Assessment of Artificial Intelligence Methods for Prediction of hERG Channel Inhibition in the "Big Data" Era. J Chem Inf Model 2020, 60 (12), 6007–6019. DOI: 10.1021/acs.jcim.0c00884 From NLM Medline.

(144) Wu, Z.; Zhu, M.; Kang, Y.; Leung, E. L.; Lei, T.; Shen, C.; Jiang, D.; Wang, Z.; Cao, D.; Hou, T. Do we need different machine learning algorithms for QSAR modeling? A comprehensive assessment of 16 machine learning algorithms on 14 QSAR data sets. Brief Bioinform 2021, 22 (4). DOI: 10.1093/bib/bbaa321 From NLM Medline.

(145) Feng, H.; Gao, K.; Chen, D.; Shen, L.; Robison, A. J.; Ellsworth, E.; Wei, G. W. Machine Learning Analysis of Cocaine Addiction Informed by DAT, SERT, and NET-Based Interactome Networks. J Chem Theory Comput 2022, 18 (4), 2703–2719. DOI: 10.1021/acs.jctc.2c00002 From NLM Medline.

(146) Zhu, Z.; Dou, B.; Cao, Y.; Jiang, J.; Zhu, Y.; Chen, D.; Feng, H.; Liu, J.; Zhang, B.; Zhou, T.; Wei, G. W. TIDAL: Topology-Inferred Drug Addiction Learning. J Chem Inf Model 2023, 63 (5), 1472–1489. DOI: 10.1021/acs.jcim.3c00046 From NLM Medline.

(147) Altae-Tran, H.; Ramsundar, B.; Pappu, A. S.; Pande, V. Low Data Drug Discovery with One-Shot Learning. ACS Cent Sci 2017, 3 (4), 283–293. DOI: 10.1021/acscentsci.6b00367 From NLM PubMed-not-MEDLINE.

(148) Shwartz-Ziv, R.; Armon, A. Tabular data: Deep learning is not all you need. Information Fusion 2022, 81, 84–90. DOI: 10.1016/j.inffus.2021.11.011.

(149) Garcia-Hernandez, C.; Fernandez, A.; Serratosa, F. Ligand-Based Virtual Screening Using Graph Edit Distance as Molecular Similarity Measure. J Chem Inf Model 2019, 59 (4), 1410–1421. DOI: 10.1021/acs.jcim.8b00820 From NLM Medline.

(150) Cereto-Massague, A.; Ojeda, M. J.; Valls, C.; Mulero, M.; Garcia-Vallve, S.; Pujadas, G. Molecular fingerprint similarity search in virtual screening. Methods 2015, 71, 58–63. DOI: 10.1016/j.ymeth.2014.08.005 From NLM Medline.

(151) Gabrielsen, M.; Kurczab, R.; Siwek, A.; Wolak, M.; Ravna, A. W.; Kristiansen, K.; Kufareva, I.; Abagyan, R.; Nowak, G.; Chilmonczyk, Z.;, et al. Identification of novel serotonin transporter compounds by virtual screening. J Chem Inf Model 2014, 54 (3), 933–943. DOI: 10.1021/ci400742s From NLM Medline.

(152) Wang, P.; Yang, F.; Yang, H.; Xu, X.; Liu, D.; Xue, W.; Zhu, F. Identification of dual active agents targeting 5-HT1A and SERT by combinatorial virtual screening methods. Biomed Mater Eng 2015, 26 Suppl 1, S2233–2239. DOI: 10.3233/BME-151529 From NLM Medline.

(153) Erol, I.; Aksoydan, B.; Kantarcioglu, I.; Salmas, R. E.; Durdagi, S. Identification of novel serotonin reuptake inhibitors targeting central and allosteric binding sites: A virtual screening and molecular dynamics simulations study. J Mol Graph Model 2017, 74, 193–202. DOI: 10.1016/j.jmgm.2017.02.001 From NLM Medline.

(154) Berfield, J. L.; Wang, L. C.; Reith, M. E. Which form of dopamine is the substrate for the human dopamine transporter: the cationic or the uncharged species? J Biol Chem 1999, 274 (8), 4876–4882. DOI: 10.1074/jbc.274.8.4876 From NLM Medline.

(155) Chen, N.; Reith, M. E. Structure and function of the dopamine transporter. Eur J Pharmacol 2000, 405 (1-3), 329–339. DOI: 10.1016/s0014-2999(00)00563-x From NLM Medline.

(156) Shelley, J. C.; Cholleti, A.; Frye, L. L.; Greenwood, J. R.; Timlin, M. R.; Uchimaya, M. Epik: a software program for pK (a) prediction and protonation state generation for drug-like molecules. J Comput Aided Mol Des 2007, 21 (12), 681–691. DOI: 10.1007/s10822-007-9133-z From NLM Medline.

(157) Pan, X.; Wang, H.; Li, C.; Zhang, J. Z. H.; Ji, C. MolGpka: A Web Server for Small Molecule pK(a) Prediction Using a Graph-Convolutional Neural Network. J Chem Inf Model 2021, 61 (7), 3159–3165. DOI: 10.1021/acs.jcim.1c00075 From NLM Medline.

(158) Neumeyer, J. L.; Tamagnan, G.; Wang, S.; Gao, Y.; Milius, R. A.; Kula, N. S.; Baldessarini, R. J. N-substituted analogs of 2 beta-carbomethoxy-3 beta- (4’-iodophenyl)tropane (beta-CIT) with selective affinity to dopamine or serotonin transporters in rat forebrain. J Med Chem 1996, 39 (2), 543–548. DOI: 10.1021/jm9505324 From NLM Medline.

(159) Bois, F.; Baldwin, R. M.; Kula, N. S.; Baldessarini, R. J.; Innis, R. B.; Tamagnan, G. Synthesis and monoamine transporter affinity of 3’-analogs of 2-beta-carbomethoxy-3-beta-(4’-iodophenyl)tropane (beta-CIT). Bioorg Med Chem Lett 2004, 14 (9), 2117–2120. DOI: 10.1016/j.bmcl.2004.02.043 From NLM Medline.

(160) Rothman, R. B.; Ananthan, S.; Partilla, J. S.; Saini, S. K.; Moukha-Chafiq, O.; Pathak, V.; Baumann, M. H. Studies of the biogenic amine transporters 15. Identification of novel allosteric dopamine transporter ligands with nanomolar potency. J Pharmacol Exp Ther 2015, 353 (3), 529–538. DOI: 10.1124/jpet.114.222299 From NLM Medline.

(161) Subbaiah, M. A. M. Triple Reuptake Inhibitors as Potential Therapeutics for Depression and Other Disorders: Design Paradigm and Developmental Challenges. J Med Chem 2018, 61 (6), 2133–2165. DOI: 10.1021/acs.jmedchem.6b01827 From NLM Medline.

(162) Glennon, R. A. The 2014 Philip S. Portoghese Medicinal Chemistry Lectureship: The "Phenylalkylaminome" with a Focus on Selected Drugs of Abuse. J Med Chem 2017, 60 (7), 2605–2628. DOI: 10.1021/acs.jmedchem.7b00085 From NLM Medline.

(163) Hsin, L. W.; Chang, L. T.; Rothman, R. B.; Dersch, C. M.; Jacobson, A. E.; Rice, K. C. Design and synthesis of 2- and 3-substituted-3-phenylpropyl analogs of 1-[2-[bis(4-fluorophenyl)methoxy]ethyl]-4-(3-phenylpropyl)piperazine and 1-[2-(diphenylmethoxy)ethyl]-4- (3-phenylpropyl)piperazine: role of amino, fluoro, hydroxyl, methoxyl, methyl, methylene, and oxo substituents on affinity for the dopamine and serotonin transporters. J Med Chem 2008, 51 (9), 2795–2806. DOI: 10.1021/jm701270n From NLM Medline.

(164) Arunotayanun, W.; Dalley, J. W.; Huang, X. P.; Setola, V.; Treble, R.; Iversen, L.; Roth, B. L.; Gibbons, S. An analysis of the synthetic tryptamines AMT and 5-MeO-DALT: emerging ‘Novel Psychoactive Drugs’. Bioorg Med Chem Lett 2013, 23 (11), 3411–3415. DOI: 10.1016/j.bmcl.2013.03.066 From NLM Medline.

(165) Brown, D. G.; Bernstein, P. R.; Wu, Y.; Urbanek, R. A.; Becker, C. W.; Throner, S. R.; Dembofsky, B. T.; Steelman, G. B.; Lazor, L. A.; Scott, C. W.;, et al. Azepines and piperidines with dual norepinephrine dopamine uptake inhibition and antidepressant activity. ACS Med Chem Lett 2013, 4 (1), 46–51. DOI: 10.1021/ml300262e From NLM PubMed-not-MEDLINE.

(166) Gruner, J. A.; Mathiasen, J. R.; Flood, D. G.; Gasior, M. Characterization of pharmacological and wake-promoting properties of the dopaminergic stimulant sydnocarb in rats. J Pharmacol Exp Ther 2011, 337 (2), 380–390. DOI: 10.1124/jpet.111.178947 From NLM Medline.

(167) Cao, J.; Slack, R. D.; Bakare, O. M.; Burzynski, C.; Rais, R.; Slusher, B. S.; Kopajtic, T.; Bonifazi, A.; Ellenberger, M. P.; Yano, H.;, et al. Novel and High Affinity 2- [(Diphenylmethyl)sulfinyl]acetamide (Modafinil) Analogues as Atypical Dopamine Transporter Inhibitors. J Med Chem 2016, 59 (23), 10676–10691. DOI: 10.1021/acs.jmedchem.6b01373 From NLM Medline.

(168) Decker, A. M.; Partilla, J. S.; Baumann, M. H.; Rothman, R. B.; Blough, B. E. The biogenic amine transporter activity of vinylogous amphetamine analogs. Medchemcomm 2016, 7 (8), 1657–1663. DOI: 10.1039/c6md00245e.

(169) Mavel, S.; Mincheva, Z.; Meheux, N.; Carcenac, Y.; Guilloteau, D.; Abarbri, M.; Emond, P. QSAR study and synthesis of new phenyltropanes as ligands of the dopamine transporter (DAT). Bioorg Med Chem 2012, 20 (4), 1388–1395. DOI: 10.1016/j.bmc.2012.01.014 From NLM Medline.

(170) Zou, M. F.; Kopajtic, T.; Katz, J. L.; Wirtz, S.; Justice, J. B., Jr.; Newman, A. H. Novel tropane-based irreversible ligands for the dopamine transporter. J Med Chem 2001, 44 (25), 4453–4461. DOI: 10.1021/jm0101904 From NLM Medline.

(171) Orjales, A.; Mosquera, R.; Toledo, A.; Pumar, M. C.; Garcia, N.; Cortizo, L.; Labeaga, L.; Innerarity, A. Syntheses and binding studies of new [(aryl)(aryloxy)methyl]piperidine derivatives and related compounds as potential antidepressant drugs with high affinity for serotonin (5-HT) and norepinephrine (NE) transporters. J Med Chem 2003, 46 (25), 5512–5532. DOI: 10.1021/jm0309349 From NLM Medline.

(172) Tosh, D. K.; Janowsky, A.; Eshleman, A. J.; Warnick, E.; Gao, Z. G.; Chen, Z.; Gizewski, E.; Auchampach, J. A.; Salvemini, D.; Jacobson, K. A. Scaffold Repurposing of Nucleosides (Adenosine Receptor Agonists): Enhanced Activity at the Human Dopamine and Norepinephrine Sodium Symporters. J Med Chem 2017, 60 (7), 3109–3123. DOI: 10.1021/acs.jmedchem.7b00141 From NLM Medline.

(173) Williard, R. L.; Middaugh, L. D.; Zhu, H. J.; Patrick, K. S. Methylphenidate and its ethanol transesterification metabolite ethylphenidate: brain disposition, monoamine transporters and motor activity. Behav Pharmacol 2007, 18 (1), 39–51. DOI: 10.1097/FBP.0b013e3280143226 From NLM Medline.

(174) Ritz, M. C.; Cone, E. J.; Kuhar, M. J. Cocaine inhibition of ligand binding at dopamine, norepinephrine and serotonin transporters: a structure-activity study. Life Sci 1990, 46 (9), 635–645. DOI: 10.1016/0024-3205(90)90132-b From NLM Medline.

(175) Froimowitz, M.; Gu, Y.; Dakin, L. A.; Nagafuji, P. M.; Kelley, C. J.; Parrish, D.; Deschamps, J. R.; Janowsky, A. Slow-onset, long-duration, alkyl analogues of methylphenidate with enhanced selectivity for the dopamine transporter. J Med Chem 2007, 50 (2), 219–232. DOI: 10.1021/jm0608614 From NLM Medline.

(176) Husbands, S. M.; Izenwasser, S.; Kopajtic, T.; Bowen, W. D.; Vilner, B. J.; Katz, J. L.; Newman, A. H. Structure-activity relationships at the monoamine transporters and sigma receptors for a novel series of 9-[3-(cis-3, 5-dimethyl-1-piperazinyl)propyl]carbazole (rimcazole) analogues. J Med Chem 1999, 42 (21), 4446–4455. DOI: 10.1021/jm9902943 From NLM Medline.

(177) Kuo, F.; Gillespie, T. A.; Kulanthaivel, P.; Lantz, R. J.; Ma, T. W.; Nelson, D. L.; Threlkeld, P. G.; Wheeler, W. J.; Yi, P.; Zmijewski, M. Synthesis and biological activity of some known and putative duloxetine metabolites. Bioorg Med Chem Lett 2004, 14 (13), 3481–3486. DOI: 10.1016/j.bmcl.2004.04.066 From NLM Medline.

(178) Lantz, K. A.; Hart, S. G.; Planey, S. L.; Roitman, M. F.; Ruiz-White, I. A.; Wolfe, H. R.; McLane, M. P. Inhibition of PTP1B by trodusquemine (MSI-1436) causes fat-specific weight loss in diet-induced obese mice. Obesity (Silver Spring) 2010, 18 (8), 1516–1523. DOI: 10.1038/oby.2009.444 From NLM Medline.

(179) Lapa, G. B.; Byrd, G. D.; Lapa, A. A.; Budygin, E. A.; Childers, S. R.; Jones, S. R.; Harp, J. J. The synthesis and biological evaluation of dopamine transporter inhibiting activity of substituted diphenylmethoxypiperidines. Bioorg Med Chem Lett 2005, 15 (22), 4915–4918. DOI: 10.1016/j.bmcl.2005.08.028 From NLM Medline.

(180) Andersen, P. H. The dopamine inhibitor GBR 12909: selectivity and molecular mechanism of action. Eur J Pharmacol 1989, 166 (3), 493–504. DOI: 10.1016/0014-2999(89)90363-4 From NLM Medline.

(181) Ananthan, S.; Saini, S. K.; Khare, R.; Clayton, S. D.; Dersch, C. M.; Rothman, R. B. Identification of a novel partial inhibitor of dopamine transporter among 4-substituted 2-phenylquinazolines. Bioorg Med Chem Lett 2002, 12 (16), 2225–2228. DOI: 10.1016/s0960-894x(02)00348-7 From NLM Medline.

(182) Han, D. D.; Gu, H. H. Comparison of the monoamine transporters from human and mouse in their sensitivities to psychostimulant drugs. BMC Pharmacol 2006, 6, 6. DOI: 10.1186/1471-2210-6-6 From NLM Medline.

(183) Solis, E., Jr.; Suyama, J. A.; Lazenka, M. F.; DeFelice, L. J.; Negus, S. S.; Blough, B. E.; Banks, M. L. Dissociable effects of the prodrug phendimetrazine and its metabolite phenmetrazine at dopamine transporters. Sci Rep 2016, 6, 31385. DOI: 10.1038/srep31385 From NLM Medline.

(184) Han, M.; Song, C.; Jeong, N.; Hahn, H. G. Exploration of 3-Aminoazetidines as Triple Reuptake Inhibitors by Bioisosteric Modification of 3-alpha-Oxyazetidine. ACS Med Chem Lett 2014, 5 (9), 999–1004. DOI: 10.1021/ml500187a From NLM PubMed-not-MEDLINE.

(185) Rothman, R. B.; Baumann, M. H.; Dersch, C. M.; Romero, D. V.; Rice, K. C.; Carroll, F. I.; Partilla, J. S. Amphetamine-type central nervous system stimulants release norepinephrine more potently than they release dopamine and serotonin. Synapse 2001, 39 (1), 32–41. DOI: 10.1002/1098-2396(20010101)39:1<32::AID-SYN5>3.0.CO;2-3 From NLM Medline.

(186) Rothman, R. B.; Vu, N.; Partilla, J. S.; Roth, B. L.; Hufeisen, S. J.; Compton-Toth, B. A.; Birkes, J.; Young, R.; Glennon, R. A. In vitro characterization of ephedrine-related stereoisomers at biogenic amine transporters and the receptorome reveals selective actions as norepinephrine transporter substrates. J Pharmacol Exp Ther 2003, 307 (1), 138–145. DOI: 10.1124/jpet.103.053975 From NLM Medline.

(187) Carroll, F. I.; Howell, L. L.; Kuhar, M. J. Pharmacotherapies for treatment of cocaine abuse: preclinical aspects. J Med Chem 1999, 42 (15), 2721–2736. DOI: 10.1021/jm9706729 From NLM Medline.

(188) Shapiro, D. A.; Renock, S.; Arrington, E.; Chiodo, L. A.; Liu, L. X.; Sibley, D. R.; Roth, B. L.; Mailman, R. Aripiprazole, a novel atypical antipsychotic drug with a unique and robust pharmacology. Neuropsychopharmacology 2003, 28 (8), 1400–1411. DOI: 10.1038/sj.npp.1300203 From NLM Medline.

(189) Efange, S. M.; Mash, D. C.; Khare, A. B.; Ouyang, Q. Modified ibogaine fragments: synthesis and preliminary pharmacological characterization of 3-ethyl-5-phenyl-1,2,3,4,5, 6-hexahydroazepino[4,5-b]benzothiophenes. J Med Chem 1998, 41 (23), 4486–4491. DOI: 10.1021/jm980156y From NLM Medline.

(190) Madras, B. K.; Xie, Z.; Lin, Z.; Jassen, A.; Panas, H.; Lynch, L.; Johnson, R.; Livni, E.; Spencer, T. J.; Bonab, A. A.;, et al. Modafinil occupies dopamine and norepinephrine transporters in vivo and modulates the transporters and trace amine activity in vitro. J Pharmacol Exp Ther 2006, 319 (2), 561–569. DOI: 10.1124/jpet.106.106583 From NLM Medline.

(191) Zhang, P.; Cyriac, G.; Kopajtic, T.; Zhao, Y.; Javitch, J. A.; Katz, J. L.; Newman, A. H. Structure-activity relationships for a novel series of citalopram (1-(3-(dimethylamino)propyl)-1- (4-fluorophenyl)-1,3-dihydroisobenzofuran-5-carbonitrile) analogues at monoamine transporters. J Med Chem 2010, 53 (16), 6112–6121. DOI: 10.1021/jm1005034 From NLM Medline.

